# Root-Suppressed Phenotype of Tomato *Rs* Mutant is Seemingly Related to Expression of Root-Meristem-Specific Sulfotransferases

**DOI:** 10.64898/2026.01.03.697460

**Authors:** Alka Kumari, Prateek Gupta, Parankusam Santisree, Injangbuanang Pamei, Satyavati Valluri, Kapil Sharma, Kavuri Venkateswara Rao, Shivani Shukla, Srilatha Nama, Yellamaraju Sreelakshmi, Rameshwar Sharma

## Abstract

Root development and growth are governed by a complex interplay between a plant’s genetic makeup and its ambient environment. In this study, we characterised a radiation-induced root-suppressed (*Rs*) mutant mapped on chromosome 4 of tomato. *Rs* seedlings had a dwarf stature, and mature plants exhibited abnormalities in their leaves, flowers, and fruits. The partial rescue of seedling root elongation by H₂S donors or L-cysteine hinted that its root defect is linked to disrupted sulfur metabolism. The metabolite profiling of organs revealed altered homeostasis, analogous to that of plants subjected to sulfur starvation. Whole-genome sequencing revealed that the *Rs* genome was replete with genetic lesions; therefore, the pleiotropic morphological abnormalities likely emanate from mutations in other genes, including those related to sulfur. Among S-related genes, two sulfotransferase genes (*Solyc04g028380* and *Solyc04g028390*) localised on chromosome 4 showed mutations in their promoters. The expression of both sulfotransferases is confined to the root meristem, suggesting a causal connection with the suppression of root growth. These sulfotransferases belonged to a distinct subset exclusively expressed in the root meristem and were distinct from the tyrosine sulfotransferase involved in the sulfation of root growth factors. Our study suggests that a sulfur-related module, other than tyrosine sulfotransferase, may regulate root development.

## Introduction

The initiation of the plant life cycle is visually marked by radicle emergence - the primary root - from germinating seeds. Post-germination, roots anchor the plant in the soil and forage for water and essential mineral nutrients. The plant’s survival depends equally on roots and shoots, the latter supplying photosynthates for root growth and function. Although radicle emergence is primarily triggered by seed hydration, subsequent root growth is governed by a complex interplay of genetic programs and environmental signals.

Molecular-genetic studies in model plants, such as *Arabidopsis thaliana,* have established that root growth is driven by sustained activity of the root meristem. The root meristem activity is modulated by the interaction between multiple genes, availability/levels of metabolites and hormones, and favorable environmental conditions. Disruption in any of the above components can block or inhibit the root growth. For instance, in Arabidopsis GRAS Transcription factor mutants, *shr* and *scr,* after radicle emergence, root growth stops, as the root meristem is unable to resume cell division (**Helariutta et al., 2000**). The root growth is also driven by the levels of plant hormones, particularly auxin, as mutants defective in auxin transport disturb auxin gradients, displaying shortened primary roots (**Blilou et al., 2005)**.

The sustenance of root growth requires shoot-derived sucrose, generated during photosynthesis. In Arabidopsis seedlings, the primary root elongation ceases after a few days of germination and is resumed only after acquisition of photoautotrophy (**Kircher and Schopfer, 2012**). The inhibition of primary root elongation in darkness is also mediated by the inactive CRY2, which associates with cell-division-promoting factors FL1 and FL3. Exposure to blue light relieves the inhibition by dissociating this complex (**Zeng et al., 2025).** The deficiency of several mineral nutrients (e.g., nitrate) inhibits primary root growth, leading to altered root architecture by initiating lateral roots (**López-Bucio et al., 2003).**

In addition to transcription factors, such as the SHR/SCR module and auxin, several peptide hormones regulate primary root development (**Hsiao and Yamada, 2020**). These peptide hormones act by controlling cell division/elongation in the root meristem. Among these peptides, the root meristem growth factors (RGFs) are expressed in the root stem cell niche. The RGFs act by binding to LRR-RLK receptors (RGF-RKs) along with SERKs as co-receptors, upregulating PLETHORA transcription factor (**Shinohara, 2021**). The upregulation of PLETHORA sustains cell proliferation in the root meristematic zone needed for continuous primary root elongation.

Post-polypeptide processing, the RGF peptides are activated by post-translational modification involving tyrosine sulfation. The sulfation of tyrosine in RGF is catalyzed by a tyrosylprotein sulfotransferase (TPST) localized in the Golgi apparatus (**Komori et al., 2009**). The TPST catalyses the transfer of sulfate from 3′-phosphoadenosine 5′-phosphosulfate (PAPS) to the side chain of tyrosine in RGF peptides. The mutation in the *TPST* gene in Arabidopsis leads to the inhibition of primary root elongation, which can be rescued by the supply of exogenous sulfated RGF peptides (**Zhou et al**., **2010)**.

In tomato and Arabidopsis, *TPST* is a single-copy gene; the loss-of-function tpst-1 Arabidopsis mutant exhibits an extremely short root phenotype. While sulfotransferase is a large gene family in plants, Arabidopsis TPST does not show any sequence similarity to other sulfotransferases (**Han et al., 2025**). Unlike TPST, the function of other plant sulfotransferases (SOTs) is not well defined; however, it is believed that these SOTs are involved in the sulfation of brassinosteroids, flavonoids, hydroxyjasmonate, and salicylic acid (**Kurogi et al., 2024).** The T-DNA knock-out mutant of *AtSOT12* (*At2g03760*) exhibited hypersensitivity to salicylic acid in seedling growth due to the loss of sulfation of salicylic acid (**Baek et al., 2010**).

In addition to the sulfation of peptides and metabolites, sulfur, via H_2_S, influences several developmental aspects of plants (**Liu et al. 2021**). In Arabidopsis, exogenous NaHS, a H_2_S donor, inhibited primary root growth by increasing the accumulation of ROS and NO (**Zhang et al. 2017**). Likewise, in tomato, a *short root* (*shr*) mutant induced by gamma rays shows inhibition of primary root elongation due to overaccumulation of NO (**Negi et al., 2010**). Similar to the *shr* mutant, the radiation-induced root-suppressed (*Rs*) mutant of tomato exhibits highly reduced root growth (**Yu and Yeager, 1960**).

We found that exogenous application of NaHS, while mildly reducing the root and hypocotyl length of control seedlings, stimulated the elongation of root and hypocotyl in *Rs* seedlings. The elongation of roots by an H_2_S donor prompted us to analyse the phenotype and metabolome to delineate the role of S-metabolism in the *Rs* phenotype. We performed whole-genome sequencing (WGS) to identify candidate genes contributing to the *Rs* phenotype. Metabolic profiling revealed that *Rs* exhibits a metabolome analogous to that of sulfur-deficient plants. WGS identified numerous mutations, including promoter-region alterations in a novel subset of root-meristem-expressed sulfotransferases localised on chromosome 4. It is plausible that these promoter mutations alter the expression of key root-specific SOTs, contributing to the severe suppression of root elongation in the *Rs* mutant.

## Results

The *Rs* mutant was initially isolated in the Chatham cultivar as a fast-neutron-induced mutant (**Yu, 1959; Yu and Yeager, 1960**). It is currently distributed as accession LA 1796 by TGRC, Davis, California, USA, in an unknown background (https://tgrc-mvc.plantsciences.ucdavis.edu/Genes/Search). In the absence of the *Rs* background information, its phenotype and biochemical characterization were carried out mainly using the Ailsa Craig (AC) cultivar as a control plant.

### A H_2_S donor stimulates the elongation of *Rs* roots

The *Rs* characteristically produces a tiny primary root, both in dark- and light-grown seedlings. Even the hypocotyls of dark- and light-grown seedlings were considerably shorter than the control cultivar AC (**Figure 1A-C**). In tomato, overproduction of NO in the roots leads to a short root phenotype, as exemplified by the *shr* mutant (**Negi et al., 2010**). However, unlike the *shr* mutant, the application of a NO scavenger cPTIO (2-(4-carboxyphenyl)-4,4,5,5-tetramethylimidazoline-1-oxyl-3-oxide) did not stimulate root elongation in the *Rs* (data not shown). Nonetheless, the application of a H_2_S donor, GYY4137 **(Li et al., 2008**), stimulated the elongation of *Rs* roots of dark-grown seedlings, while it had little effect on AC (**Figure 1C-D).** A similar stimulation of *Rs* root elongation, albeit milder, was elicited by sulfide donor NaHS (**Whiteman et al., 2010**) in a dose-dependent manner up to 100 µM, beyond which stimulation declined. Contrary to GYY4137, NaHS also elicited the hypocotyl elongation of the *Rs* seedlings, while NaHS-induced stimulations were lacking in AC (**Figure 1E; Figure S1A).**

**Figure 1:**
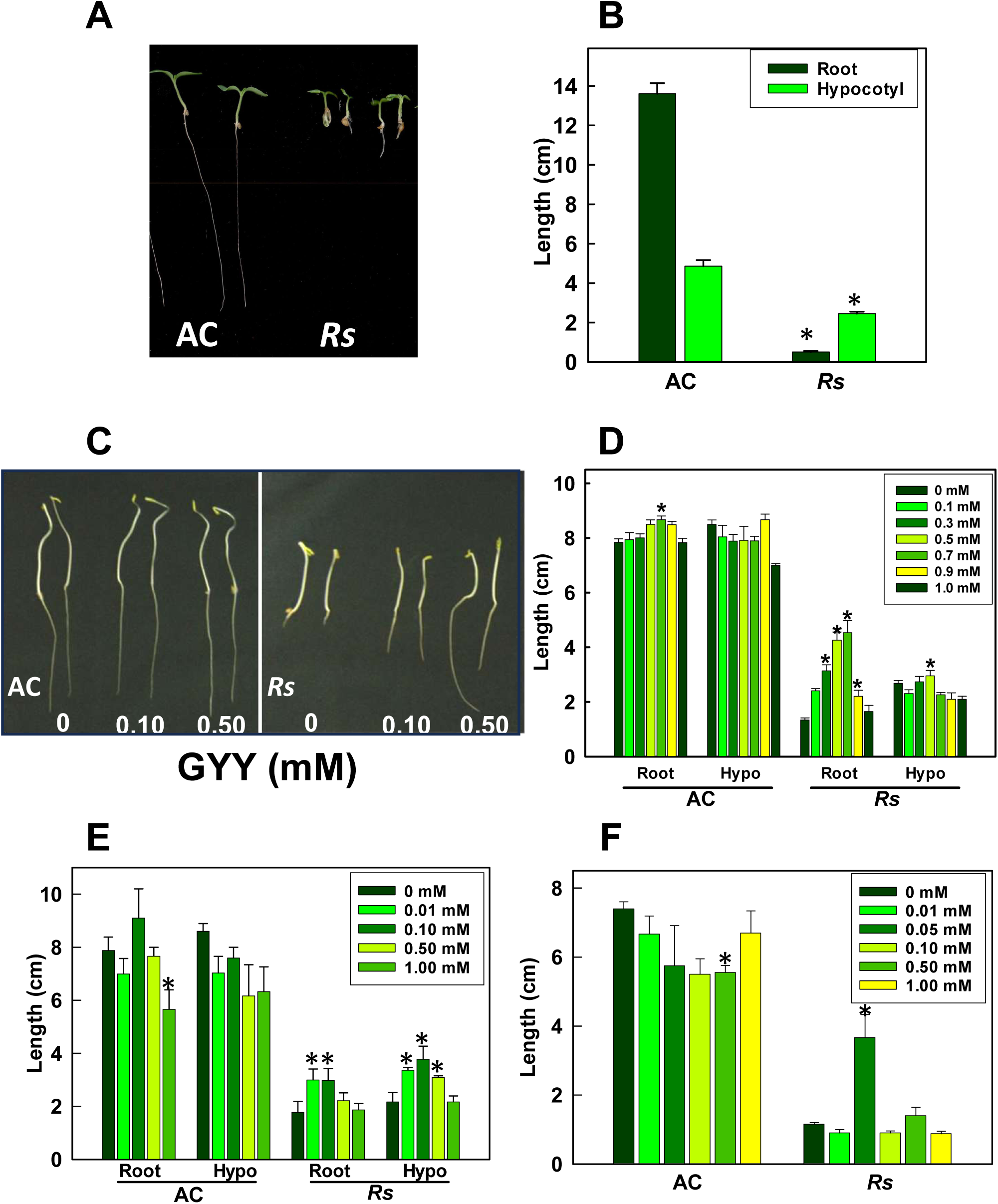
Influence of H_2_S donors on the *Rs* mutant root. **A-B.** Light-grown seedlings (5-day-old) of the *Rs* mutant showing shorter hypocotyl and root compared to Ailsa Craig. **C-D.** Dark-grown seedlings (5 days old) of the *Rs* mutant exhibit a **s**uppressed root phenotype. The H_2_S donor GYY4137 stimulates root elongation in *Rs* in a concentration-dependent manner up to 0.7 mM. GYY4137 does not affect the hypocotyl length in *Rs* and the hypocotyl/root length in AC. **E.** The H_2_S donor NaHS also stimulates root elongation in *Rs* seedlings. **F.** The Sulfur-containing amino acid L-cysteine stimulates root elongation in *Rs,* while D-cysteine is ineffective (**See Figure S1**). The asterisk (*) indicates a statistically significant difference between treated and control plants (P<0.05).

Considering that endogenous H_2_S is produced by enzymatic desulfuration of L-cysteine, we examined whether L-cysteine can stimulate the primary root elongation of *Rs* (**Figure 1F**). The stimulation of *Rs* root elongation by L-cysteine was restricted to a narrow window of 0.05 mM concentration, while higher dosages reduced the stimulation. The stimulation of root elongation was restricted to L-cysteine; D-cysteine at a similar concentration did not elicit any stimulation of root elongation (**Figure S1B**). To further confirm the role of H_2_S in root elongation, we investigated whether a H_2_S scavenger, hypotaurine (**Regelmann and Grieshaber, 2008**), would reduce the root elongation of AC seedlings. The dosage-dependent reduction in the root length of AC seedlings treated with hypotaurine indicated that the removal of H_2_S impedes the elongation of the roots (**Figure S1C**). Likewise, the application of DL-propargylglycine (PAG, an inhibitor of H_2_S biosynthesis (**Kaya et al., 2020**)) also drastically reduced both root and hypocotyl elongation in the AC and *Rs* seedlings (**Figure 2A, Figure S1D**).

**Figure 2:**
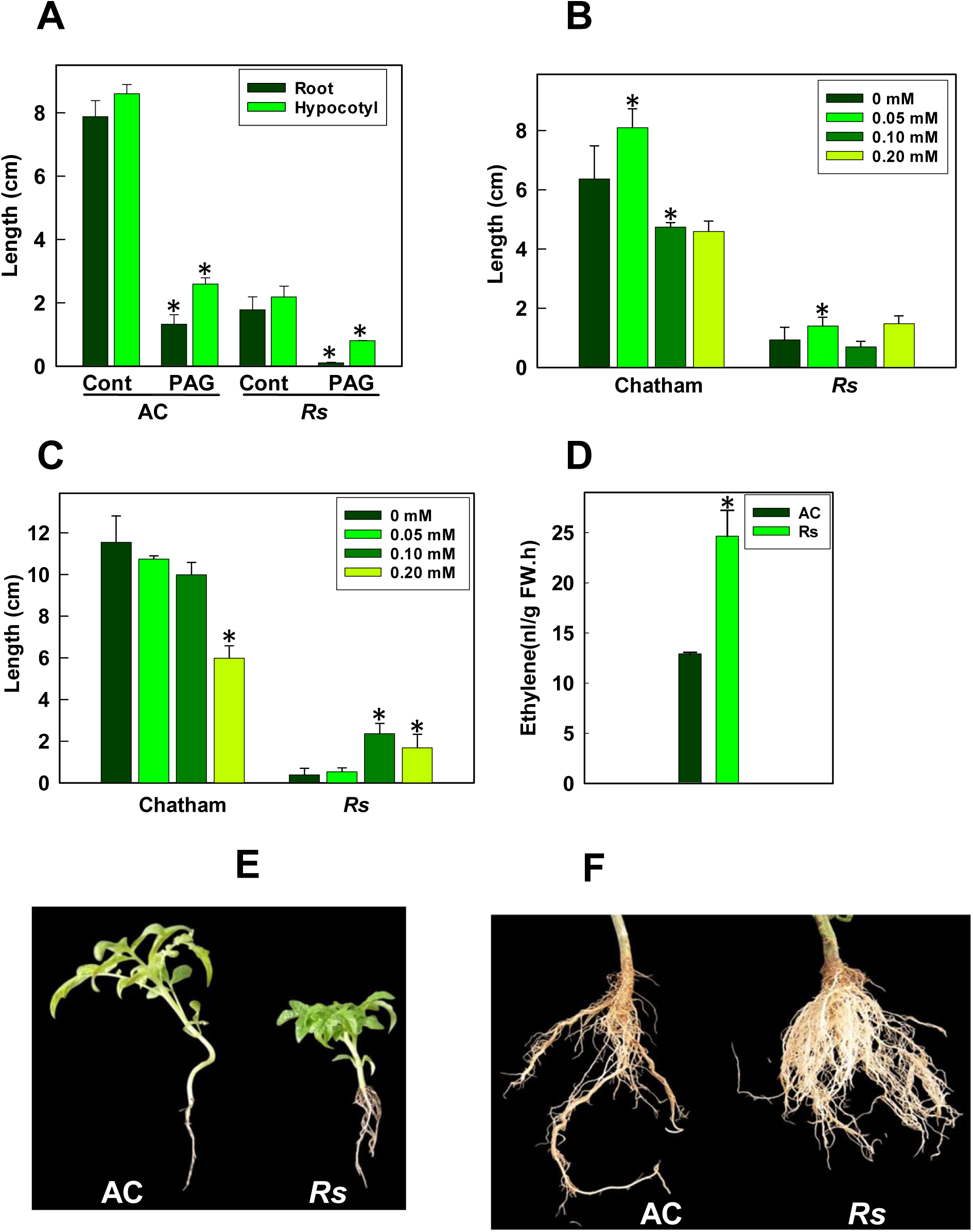
Characterization of *Rs* mutant response and phenotype. **A.** The dark-grown seedlings (7-day-old) treated with DL-propargylglycine (PAG), an inhibitor of cystathionine γ-lyase (CSE), show strong inhibition of hypocotyl and root elongation in *Rs* and AC seedlings **B-C.** Comparison of L-cysteine-stimulated root elongation in dark-grown (**B**) and light-grown (**C**) *Rs* seedlings (5-day-old) with its progenitor cultivar Chatham as control. **D**. Ethylene evolution from dark-grown seedlings of *AC* and *Rs* mutants. Phenotype of three-week-old greenhouse-grown young AC and *Rs* plants. **E.** Mature roots collected from three-month-old greenhouse-grown AC and *Rs* plants. Note, Post-seedlings stage, *Rs* displays a more branched root phenotype by initiating more lateral roots (**D-E**). The asterisk (*) indicates a statistically significant difference between treated and control plants (P<0.05).

Since the *Rs* mutant was initially isolated in the Chatham background, we examined whether Chatham displays root elongation in the presence of L-cysteine. As expected, a 0.05 mM concentration of L-cysteine stimulated the root elongation in dark-grown *Rs* seedlings. At higher concentrations, L-cysteine suppressed the root elongation in dark-grown Chatham seedlings. Similar inhibition of root elongation at higher L-cysteine concentration was also observed for light-grown Chatham seedlings. Like dark-grown seedlings, the light-grown *Rs* seedlings also showed root elongation, albeit at 0.1 mM and above concentrations of L-cysteine (**Figure 2B-C; Figure S2**).

Since dark-grown *Rs* seedlings show shorter roots and hypocotyls than AC, the *Rs* phenotype is similar to the *Epi* mutant (**Barry et al., 2001**), which, due to ethylene overproduction, has stunted seedlings. We examined ethylene emission from dark-grown *Rs* seedlings to ascertain whether the *Rs* mutation affected ethylene biosynthesis. Compared to AC, the *Rs* seedlings emitted nearly 2-fold higher ethylene (**Figure 2D**). To determine whether the *Rs* phenotype was related to ethylene, we compared the effect of GYY4137 on root elongation in the *Epi* mutant. We also included the *Nr* mutant, an ethylene-insensitive mutant, due to a mutation in the ethylene receptor ETR3 (**Wilkinson et al., 1995**). Unlike *Rs* seedlings, GYY4137-treated *Epi* and *Nr* mutant seedlings did not elicit root elongation (**Figure S3A**). *Epi* and *Nr* mutants showed only mild modulation of their root length, while *Rs* roots elongated significantly. Ostensibly, the *Rs* mutant phenotype is due to a lesion other than ethylene overproduction. In Sweet Pepper, the salinity-induced inhibition of root growth can be mitigated by applying glutathione and glutathione plus NaHS, which induces H_2_S emission (**Kaya et al., 2020**). In contrast to this, glutathione application inhibited the root elongation in AC, *Nr*, *Epi,* and *Rs* seedlings; thus, discounting any role of glutathione in the stimulation of root elongation in *Rs* seedlings (**Figure S3B**).

### *Rs* plants displayed abnormal development

The restricted root growth phenotype of the *Rs* mutant was observed only during the early seedling stage. During the vegetative growth, post-seedling stage, it generated more lateral roots, thus compensating for restricted primary root development (**Figure 2E**). At the end of the life cycle, the *Rs* plants have more branched root biomass than that of AC (**Figure 2F**). Compared to AC and Chatham, the *Rs* plants were of shorter stature (**Figure 3A**). The leaves of the *Rs* plants were more compact, highly serrated, and had more chlorophyll than those of AC (**Figure 3B-C**). The reproductive phenotype of the *Rs* plant also differed from that of AC (**Figure 3D-E**). A prominent feature was the more compact inflorescence displayed by *Rs* compared to AC. The *Rs* plants had twice as many inflorescences as AC; however, the number of flowers per inflorescence was lower than that of AC.

**Figure 3.**
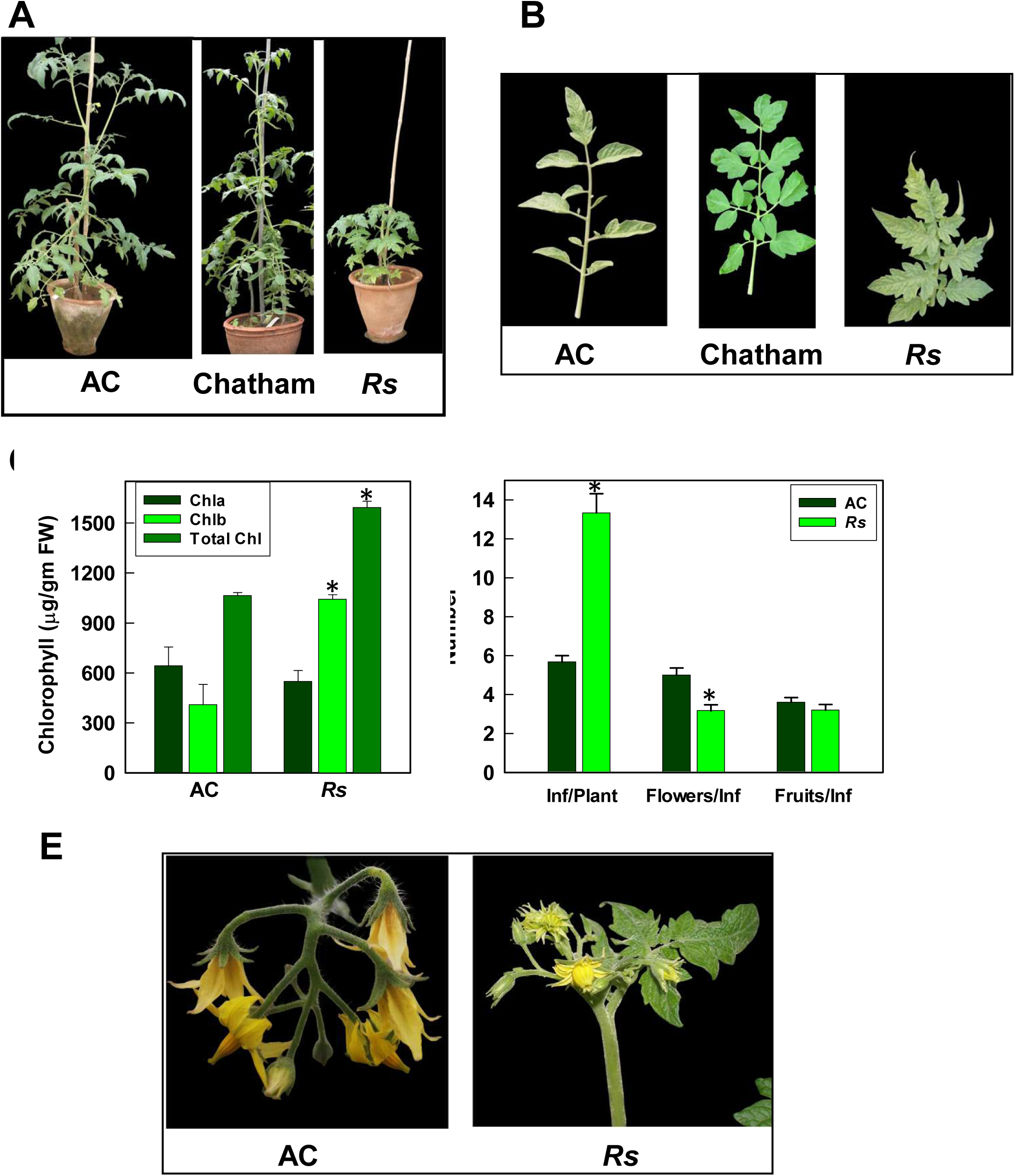
Root-suppressed (*Rs*) mutant exhibits stunted plant growth. **A.** Phenotypes of tomato AC, Chatham, and Rs plants. **B.** Leaf of AC, Chatham, and *Rs* from the eighth node of two-month-old plants. Note the variation in leaf size, shape, and color in the *Rs* mutant. **C.** Quantification of leaf chlorophyll content. **D.** Comparison of flowers, inflorescences, and fruit numbers of AC and *Rs* plants. **E.** Inflorescence of AC and *Rs* plants. The asterisk (*) indicates a statistically significant difference between treated and control plants (P<0.05).

Some flowers of *Rs* displayed abnormal development, characterised by an increased number of sepals and petals, and an open anther cone (**Figure 4A-B**). The fruits arising from the abnormally shaped flower had a convoluted shape (**Figure 4C**). The mature green fruits of the *Rs* mutant were lighter green than those of AC (**Figure 4D**). The shape of red-ripe fruits varied widely, and this variation in shape was linked to abnormalities in floral morphology. Overall, the *Rs* fruits were larger than those from Chatham. The red ripe fruits of the *Rs* had a pinkish hue on the skin. (**Figure 4E**). Consistent with this, the anthocyanin level in *Rs* fruits was three times higher than in Chatham (**Figure S4A**). The *Rs* fruits had a slightly larger horizontal perimeter than those of Chatham (**Figure S4B**). The cut section of the *Rs* fruits revealed that the fruits had a thicker septum, while the abnormally shaped fruits had an elongated septum (**Figure 4F-G**). The total carotenoid level and the distribution of carotenoids in *Rs* were nearly identical to those in Chatham (**Figure 4H, Dataset S1**).

**Figure 4.**
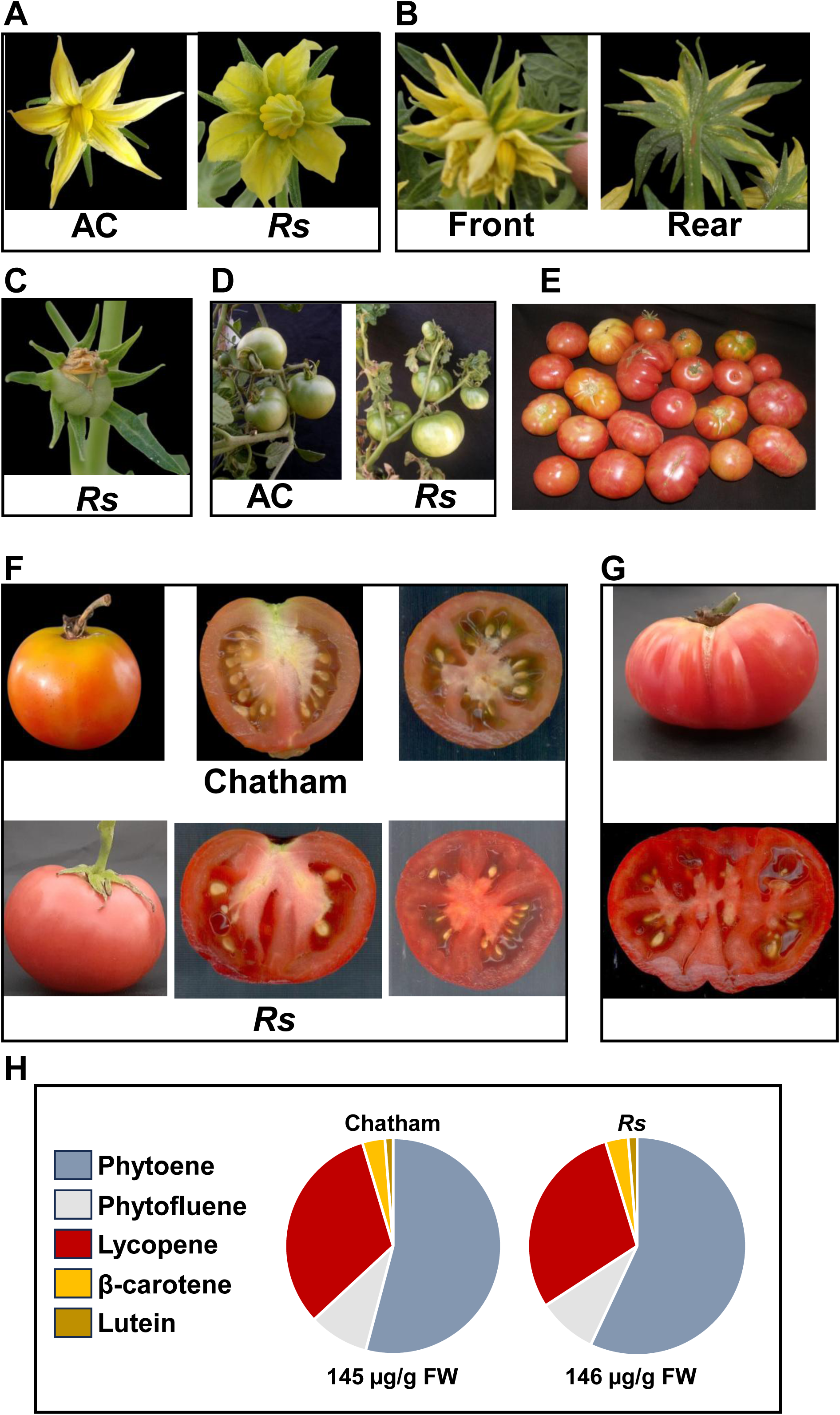
The *Rs* flowers and fruits compared to controls**. A.** AC and *Rs* flower. Note the wide petals and a large anther cone in the *Rs* flower. **B**. Abnormal flower of *Rs* with multiple petals and twisted shape. Left- Front view; Right- Rear view. **C**. A fruit developing from an abnormal flower. **D.** Green fruits of AC and *Rs* plants. Mild green stripes are visible near the pedicle. **E.** The *Rs* fruits show variation in shape. The fruits have a pink skin texture. **F.** Internal structure of Chatham and *Rs* fruits. Left- intact fruit, Middle-Vertically sliced half, Right- Horizontally sliced fruit half. **G**. Abnormal *Rs* fruit (top) and its horizontally sliced half (Bottom). **H**. Relative contents of different carotenoids in red-ripe fruits of Chatham, and *Rs*. The values below each pie indicate the total carotenoid content (µg/g fresh weight). Note that the area of the pie is proportional to the total amount of carotenoids. See **Dataset S1** for individual carotenoid levels and significance. The carotenoid data are expressed as mean SE (n ≥ 3; P ≤ 0.05).

### A massive metabolic shift was observed in the *Rs* seedling

The partial rescue of the root phenotype by sulphur-donating molecules suggested the possibility of metabolic alteration in the *Rs* mutant. To investigate this, we compared the metabolomes of the *Rs* mutant and AC across developmental stages (seedlings, roots, leaves, and floral buds). The metabolome of the fruits of the *Rs* mutant was compared with that of Chatham, as the *Rs* mutant was first isolated in Chatham and later introgressed into other cultivars (**Yu, 1959)**.

Principal Component Analysis (PCA) revealed a unique and distinct metabolomic profile of *Rs* seedlings compared to AC, showing close clustering within *Rs* and clear separation from AC (**Figure 5A-C, Dataset S2**). Compared to seedlings, the mature roots of *Rs* and AC grouped more closely, suggesting a broadly similar metabolome with minor differences. At the same time, the metabolome of mature roots is widely different from that of seedlings. The PCA of the 4th node leaf of *Rs* also showed clear separation from the AC, whereas the 8th node leaf of the AC and the *Rs* showed more similarity. The substantial overlap of PCAs of floral buds indicated high metabolic similarity between *Rs* and AC flower buds. In contrast, the PCA of Chatham and *Rs* fruits was separated, reflecting that while the metabolomes of *Rs* and Chatham fruits are distinct, there is more heterogeneity in their respective metabolome.

**Figure 5.**
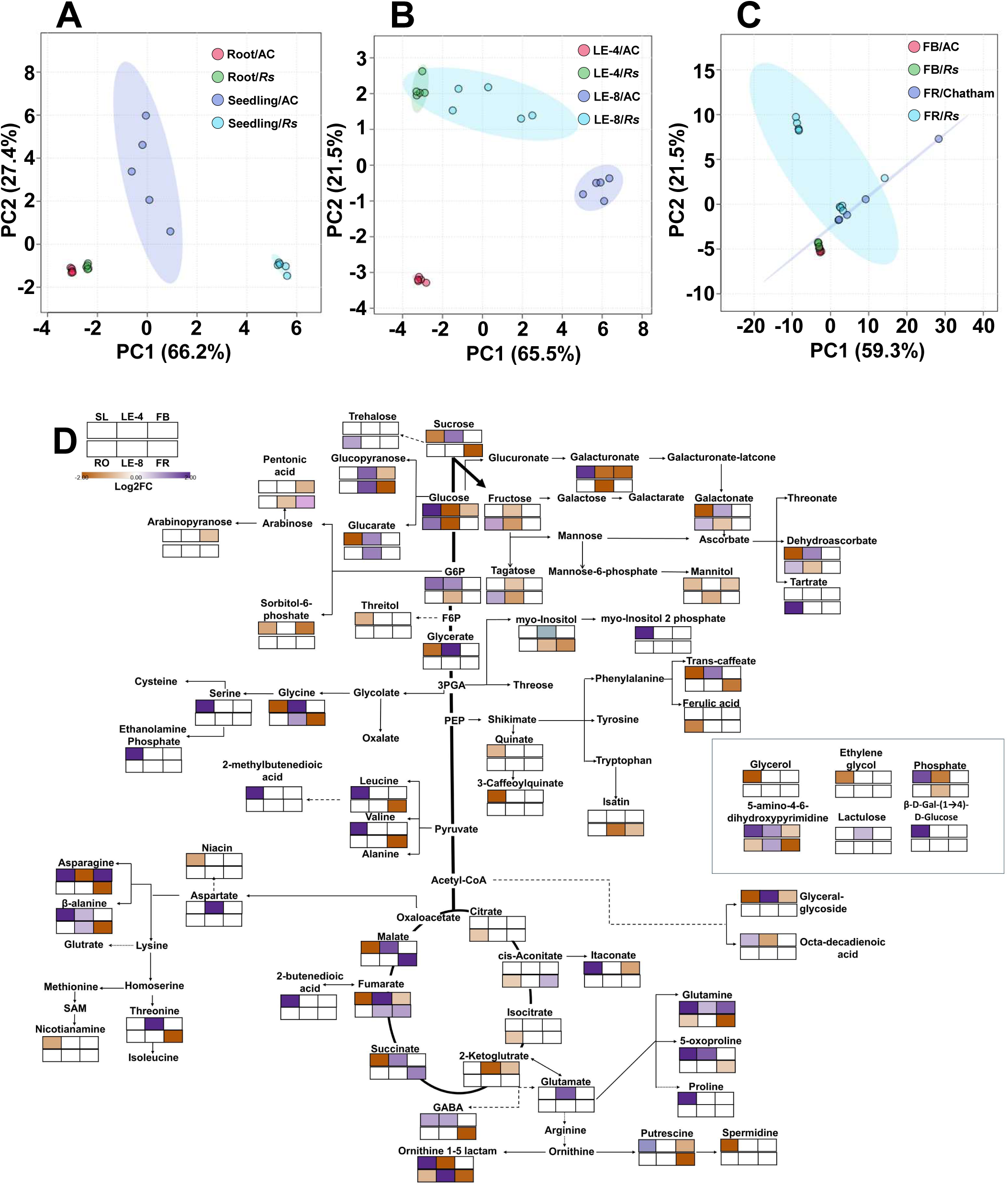
Metabolite profiles of different organs of the *Rs* mutant. (**A-C**) Principal component analysis (PCA) of metabolites. **A-** Seedlings and Root. **B-** 4^th^ node Leaf and 8th node leaf. **C-** Floral buds and fruits. The PCA was constructed using MetaboAnalyst 6.0. The variances of the PC1 and PC2 are given within parentheses. (**D**) The levels of metabolites in different organs of *Rs* nutant relative to AV or Chatham. The relative changes in metabolite levels were calculated as the *Rs*/AV ratio for each organ, barring fruit, where Chatham was used as a control. The grid in the top left corner of the figure represents the organs used. Only metabolites with significant changes (log2 fold change ≥ ± 0.584; P-value ≤ 0.05; n ≥ 3) are displayed on the metabolic pathway. For detailed metabolite data, refer to **Datasets S2** and **S4.** Abbreviations. **SL**- Seedlings; **RO**-Mature roots; **LE-4**- Leaf from the 4^th^ node of the plant; **LE-8**- Leaf from the 8th node of the plant; **FB**- Floral bud; **FR**- Red ripe fruits.

Consistent with PCA, the metabolome of the *Rs* displayed organ-specific variation in metabolite levels. Of the 72 detected metabolites, 62 varied in the *Rs* mutant at one or more developmental stages. A six-set Venn analysis indicated only a few overlapping up- or down-regulated metabolites between different developmental stages (**Figure S5, Dataset S3**). The most substantial changes occurred in *Rs* seedlings, where 23 metabolites were upregulated (↑) and 21 were downregulated (↓) (**Dataset S4**). In contrast, in *Rs* fruits, most metabolites were downregulated (20↓), and only a few were upregulated (3↑). Floral buds and mature roots showed the lowest variability with the least number of up- and down-regulated metabolites (Floral bud- 2↑, 14↓; Mature root- 8↑, 7↓). The 4^th^ node and 8^th^ node leaves showed a contrasting pattern- the 4th node leaf had more upregulated metabolites (23↑, 9↓), while the 8th node leaf had more downregulated metabolites (6↑, 14↓). Several metabolites in leaves displayed shared regulation patterns (5↓, 4↑) while a few (6) exhibited opposite regulation.

### Branched-chain amino acids were upregulated in *Rs* Seedlings

In *Rs* seedlings, significant upregulation of branched-chain amino acids (valine, leucine) and nitrogen-rich compounds (glutamine, asparagine) indicated significant changes in amino acid metabolism (**Figure 5D; Figure S6**). A marked downregulation of glycine accompanied the above upregulation. Additionally, the downregulation of succinic acid and dehydroascorbic acid, alongside increased levels of 2-butenedioic and itaconic acids, pointed to altered organic acid metabolism. Enhanced glycolysis was suggested by increased glucose and glucose-6-phosphate. Alterations in myo-inositol-2-phosphate, phenolics, and other metabolites further supported a substantial metabolic reprogramming in *Rs* seedlings.

In contrast, *Rs* fruits showed reduced branched-chain amino acids (valine, leucine), stress-associated proline, and nitrogen-rich compounds (glutamine, asparagine). A significant increase in malic acid indicated a shift in the TCA cycle toward acid accumulation during the ripening process. Downregulation of β-D-glucopyranose and sucrose indicated altered carbohydrate metabolism. Reduced levels of myo-inositol, t-caffeic acid, and putrescine suggested decreased stress signalling in *Rs* fruits.

In contrast to seedlings, the mature *Rs* root displayed minimal change in amino acid metabolism, with only β-alanine (↑) and glutamine (↓) altered. The upregulation of tartaric acid and downregulation of other TCA cycle intermediates (citric, isocitric, and cis-aconitic acids) indicated a shift towards tartrate accumulation. Increased glucose levels, fructose, trehalose, and tagatose suggested enhanced sugar accumulation or energy storage. These subtle changes were consistent with the PCA proximity between *Rs* and AC roots.

In the *Rs* 4th node leaf, upregulation of TCA cycle intermediates (Fumaric, Malic, Succinic acids), except for the downregulation of 2-ketoglutaric acid, indicated an upregulation of TCA metabolism-linked processes. Likewise, the upregulation of amino acids (glycine and aspartic acid) and sugars (sucrose and β-D-glucopyranose) indicates a shift in carbon allocation for export to the sink. Compared to the 4th node leaf, fewer differentially regulated metabolites were detected in the 8^th^ leaf. Moreover, most metabolites were downregulated in the 8th node leaf, reflecting diversity in individual leaf metabolomes. Between the 4^th^ and 8^th^ node leaves, eleven metabolites were similarly up- or down-regulated. The downregulation of glucose, fructose, and tagatose indicated a shift toward the formation of monosaccharides in the 4th and 8th node leaves.

*Rs* floral buds exhibited minimal deviation from AC, like the *Rs* root, with few differentially expressed metabolites. Upregulation of asparagine and glutamine indicated enhanced nitrogen metabolism in *Rs* buds. The downregulation of TCA cycle intermediates (2-Ketoglutaric, Fumaric acids), associated organic acids (2-butenedioic, Itaconic acids), and glucose indicated reduced metabolism in *Rs* floral buds. The metabolome of *Rs* floral buds was predominantly downregulated, indicating lowered metabolism relative to AC.

### Impaired sulfotransferase expression may be the genetic basis for the *Rs* phenotype

The partial recovery of root growth in the *Rs* mutant by H₂S and L-cysteine suggested a defect in sulfur metabolism or signalling. Classical morphological mapping had previously localised the *Rs* mutation to chromosome 4 (**Kerr, 1972**), using the *entire* mutant (now identified as ***Solyc04g076850*** by **Wang et al., 2005**) as a morphological marker. Whole-genome sequencing of the *Rs* mutant and comparison with the Heinz reference genome identified mutations in several sulfur-related genes (**Dataset S5**). In chromosome 4, three genes were identified related to S-metabolism and signal transmission: two sulfotransferases (***Solyc04g028380*, *Solyc04g028390***) and a RGF receptor kinase (RGF-RK) (***Solyc04g064940***). All three genes had mutations in their promoter region. Among these genes, a sulfotransferase (***Solyc04g028390***) was the most promising candidate, as it contained three insertions and a transversion in its promoter region (**Dataset S5; Table S1**). These insertions deleted the binding sites of several transcription factors and, in parallel, added binding sites for additional TFs (**Dataset S6; Table S2).** Given that *Solyc04g028390* expression is solely localised to the root meristem (**Figure 6**; https://conekt.sbs.ntu.edu.sg) (**Proost and Mutwil, 2018**), we infer that these mutations alter its expression, resulting in the root-suppressed phenotype.

**Figure 6.**
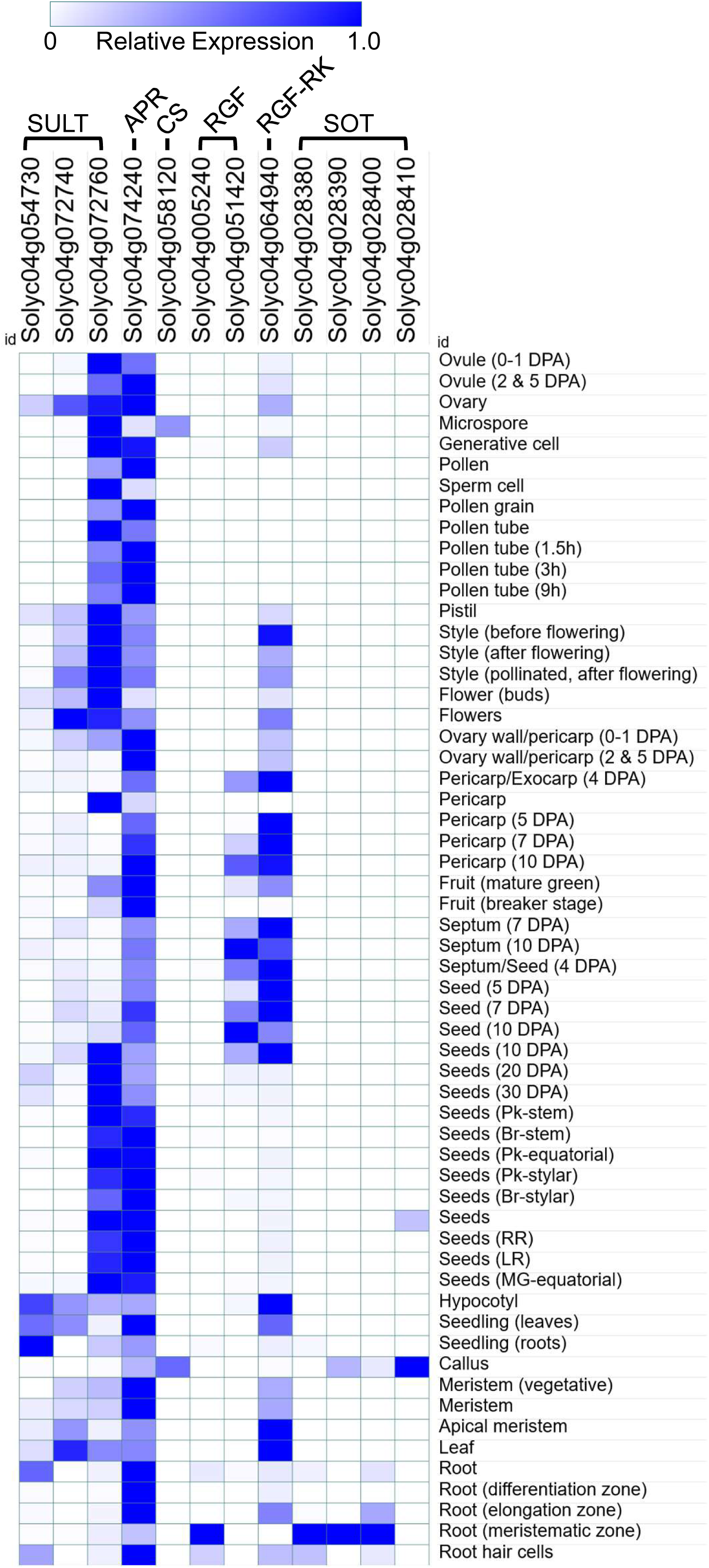
Tissue-specific expression profile of sulfur metabolism/signalling-related genes localised on chromosome 4 of tomato. The depicted genes from left to right are Sulfate transporters (**SULT**, Solyc04g054730, Solyc04g072740, Solyc04g072760); 5’-adenylylsulfate reductase (**APR**, Solyc04g074240); Cysteine synthase (**CS**, Solyc04g058120); Root growth promoting factor (**RGF**, Solyc04g005240, Solyc04g 051420), RGF receptor kinase (**RGF-RK**, Solyc04g064940) and Sulfotransferase (**SOT**, Solyc04g028380, Solyc04g028390, Solyc04g028400, Solyc04g028410). Mean expression data were sourced from the Conekt platform (https://conekt.sbs.ntu.edu.sg). Expression values for each gene were normalized to their maximum expression level observed in any tissue (set to 1). The heatmap displays the relative expression level of each gene across various tissues. The accompanying color scale bar indicates the relative expression intensity. ***Note:*** Three sulfotransferase genes (Solyc04g028380, Solyc04g028390, and Solyc04g028400) and one RGF gene (Solyc04g005240) show prominent expression in the root meristem. Among these, two sulfotransferases (Solyc04g028380, Solyc04g028390) and RGF receptor kinase (RGF-RK, Solyc04g064940) have mutations in the promoter. For the relative values for each gene, see **Dataset S7**.

The shortening of the root can also result from mutations in RGF peptides or their receptor, the RGF-RK. Though RGF, *Solyc04g005240*, is also exclusively expressed in the root, it (*Solyc04g005240*) and another RGF (*Solyc04g051420*) had no mutations either in the gene or the promoter. On the other hand, RGF-RK, *Solyc04g064940*, has an insertion in the promoter altering transcription factor binding sites. However, unlike the root-meristem-specific expression of *Solyc04g028390* sulfotransferase, RGF-RK (*Solyc04g064940*) is expressed in all organs.

In Arabidopsis, RGFs are activated by tyrosine sulfation, a reaction catalysed by the tyrosine sulfotransferase AtTPST (AT1G08030). Mutations in *AtTPST* result in a stunted primary root, a phenotype reminiscent of the *Rs* mutant. However, Solyc04g028390 protein shares no homology with AtTPST, indicating it belongs to a distinct sulfotransferase group. Phylogenetic analysis places the Solyc04g028390 protein within a unique clade of four sulfotransferases on tomato chromosome 4 (**Group VI, Figure S7; Faraji et al., 2020**). Notably, three of these four sulfotransferase genes, including *Solyc04g028390* and a RGF, *Solyc04g005240*, exhibit predominant expression in the root meristem, reinforcing their potential role in root development (**Figure 6**).

While the root suppression in *Rs* is likely attributable to the altered expression of *Solyc04g028390*, the pleiotropic vegetative and reproductive phenotypes, as well as the broader shifts in sulfur metabolism, may stem from mutations in other genes in the *Rs* genome. To study the influence of all S-related mutated genes in the *Rs* genome on metabolism, we examined their expression in various organs of tomato. (**Figure 7; Dataset S7)** shows the expression of sulfur-metabolism-related genes in different organs of tomato sourced from the Conekt platform (https://conekt.sbs.ntu.edu.sg). Expectedly, the mutations in the above genes in *Rs* may alter their expression and, in turn, influence metabolic homeostasis. The adenylyl-sulfate kinase (***Solyc10g080680***) that provides PAPS for sulfation of RGFs is expressed in all organs of tomato. The mutated *Solyc10g080680* gene in *Rs* may reduce the availability of PAPS, thus exacerbating the *Rs* phenotype. Likewise, the PAPS reductase gene (***Solyc03g031620***) is maximally expressed in the floral buds. In *Rs*, the mutation in *Solyc03g031620* could impair its function and contribute to the observed metabolic alterations in floral buds. Likewise, mutations in other sulfur-related genes that are highly expressed in fruit tissues may be responsible for the altered metabolic homeostasis of *Rs* fruits. To sum, the wide range of metabolic and morphological changes in *Rs* reflects the combined effect of the mutations present across its genome.

**Figure 7.**
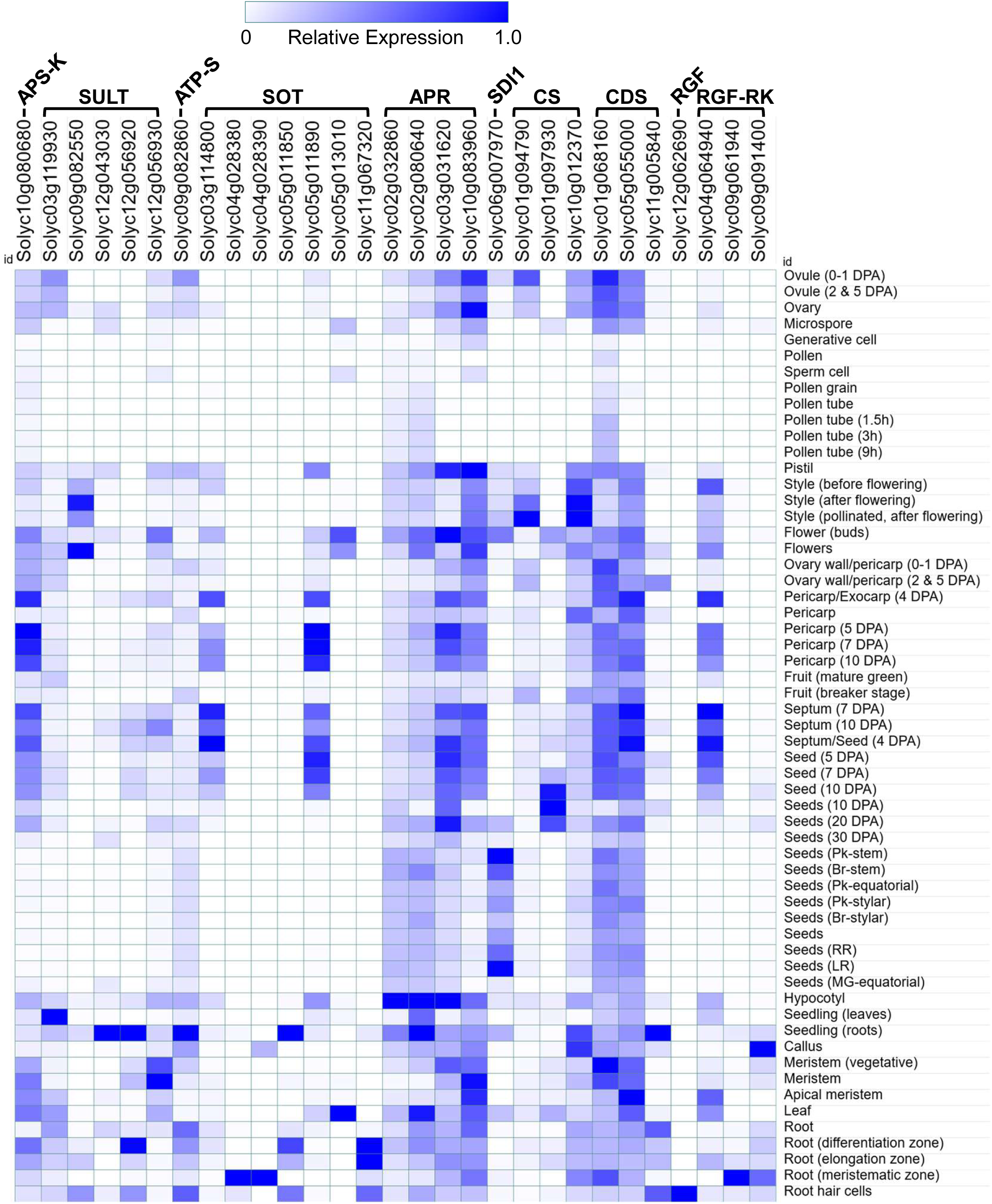
Tissue-specific expression profile of S-metabolism-related genes in tomato. The heatmap displays the relative expression levels of only those genes in which mutations were detected in the *Rs* genome. It is expected that the mutations may shift the above-observed expression patterns, in turn, altering S-metabolism in the *Rs* mutant. The genes displayed are from left to right, are adenylate-sulfate kinase (**APS-K**, Solyc10g080680), Sulfate transporter (**SULT**- Solyc03g119930, Solyc09g082550, Solyc12g043030, Solyc12g056920, Solyc12g056930), ATP sulfurylase (**ATP-S**, Solyc09g082860), Sulfotransferase (**SOT,** Solyc03g114800, Solyc04g028380, Solyc04g028390, Solyc05g011850, Solyc05g011890, Solyc05g013010, Solyc11g067320), 5’-Adenylylsulfate reductase (**APR**, Solyc02g032860, Solyc02g080640, Solyc03g031620, Solyc10g083960), Sulfur Deficiency-induced 1 (**SDI1**, Solyc06g007970), Cysteine Synthase (**CS**, Solyc01g094790, Solyc01g097930, Solyc10g012370), Cysteine desulfurase (**CDS**, Solyc01g068160, Solyc05g055000, Solyc11g005840), Root growth factor (**RGF**, Solyc12g062690) and RGF receptor kinase (**RGF-RK,** Solyc04g064940, Solyc09g061940, Solyc09g091400). **Note:** Among these genes, Solyc04g028380 and Solyc04g028390 are exclusively expressed in the root meristem. Mean expression data were sourced from the Conekt platform (https://conekt.sbs.ntu.edu.sg). Expression values for each gene were normalized to their maximum expression level observed in any tissue (set to 1). The accompanying color scale bar indicates the relative expression intensity. For the relative values for each gene**, see Dataset S7.**

## Discussion

Establishing a direct genotype-phenotype relationship for the *Rs* mutation is difficult due to the presence of three secondary mutations, which may also influence the observed phenotypes (**Table S3**). These additional mutations may have acted independently or interdependently along with the *Rs* mutation during the life course of *Rs* plants. It is plausible that the dwarf and green-leaves phenotype of *Rs* is related to the *d* (dwarf-Brassinosteroid C-6 Oxidase) mutation (**Nomura et al., 2005**), which is part of the LA1796 accession. However, the *d* mutant of tomato does not exhibit a short root phenotype in dark-grown seedlings, at least in the Micro-tom background, although it has short hypocotyls, similar to the *Rs* mutant (**Martí et al., 2006**). The LA1796 accession also harbours *gs* (green stripe-Methylation of the *TAGL1* promoter) (**Liu et al., 2020**) and *h* (hair absent- C2H2-type domain-containing protein) (**Chang et al., 2018**) mutations (**Table S3**). Consistent with harbouring the *h* mutation, *Rs* plants had fewer hairs on the vegetative and reproductive organs than AC (**Figure 3E**). Likewise, the green fruits of the *Rs* mutant also showed mild green stripes near the pedicel region (**Figure 4D**). However, the *h* and *gs* mutation phenotypes are restricted to their respective organs, and no adverse effects on seedling phenotypes have been reported in the literature. The *Rs* being a dominant mutant is also genetically distinct from *d, h,* and *gs* mutants, which are recessive.

### H_2_S Donors Stimulate *Rs* Root Elongation

The *Rs* mutant isolated in the Chatham background had extremely suppressed roots, or no roots at all. However, the seedlings of the LA1796 accession largely displayed short roots, and only occasional seedlings had a complete root-suppressed phenotype. This may be related to the expressivity of the *Rs* mutation in the unknown background. A distinct feature was that *Rs* exhibited short roots both in dark- and light-grown seedlings, and typical light-stimulated primary root elongation was absent in *Rs* seedlings (**Al-Hammadi et al., 2003**). The rescue of mutant phenotype by the external application of chemicals can give a clue about the metabolic lesion underlying the mutation. The short root phenotype of the tomato *shr* mutant can be rescued by scavenging of NO, hinting that hyperaccumulation of NO leads to shortening of the root (**Negi et al., 2010**). Similarly, the stimulation of *Rs* root elongation by H_2_S donors, such as (GYY4137 and NaHS), suggests a likely block in sulphur assimilation/transport/signalling is responsible for the shortening of the root.

The differential efficacy of GYY4137 and NaHS in stimulating *Rs* root elongation can be attributed to their distinct H_2_S release kinetics. While GYY4137 releases H_2_S in a slow and sustained manner over a longer period, the release of H_2_S from NaHS is transient, contributing to observed variation. Hypotaurine is a potent H_2_S scavenger (**Ortega et al., 2008**) and has been utilised in plants to reduce intracellular H_2_S content (**Li et al., 2014**). The shortening of AC roots by hypotaurine treatment indicates that H_2_S is needed for normal root elongation, even in control seedlings.

### L-Cysteine too stimulates *Rs* Root Elongation

The stimulation of *Rs* root elongation by H_2_S suggests a likely defect within the sulphur assimilation pathway, either in transport or in the conversion of inorganic sulphate (SO_4_^2-^) to reduced sulphur forms used for synthesising essential sulphur-containing amino acids like cysteine. The stimulation of *Rs* root elongation by L-cysteine but not by D-cysteine indicates that the reduced synthesis or availability of L-cysteine may have contributed to the shortening of *Rs* roots. In plants, cysteine desulfhydrase degrades cysteine into H_2_S, ammonia, and pyruvate (**Papenbrock et al., 2007**), which may have generated H_2_S in L-cysteine-treated seedlings. Consistent with this, the knockout mutants of L-cysteine desulfhydrase reduce endogenous H_2_S production and trigger premature leaf senescence in tomato (**Hu et al., 2021**) DL-propargylglycine acts as an inhibitor of cysteine desulfhydrases (**Kaya et al., 2020**). Considering that DL-propargylglycine strongly inhibits both hypocotyl and root elongation, it suggests that a block in the formation of cysteine and H_2_S may have contributed to the *Rs* root phenotype. The rescue of the *Rs* root phenotype seems to be limited to L-cysteine, as exogenous glutathione did not stimulate root elongation. The *Rs* seedlings emitted higher ethylene levels, similar to the *Epi* mutant, an overproducer of ethylene, which also displayed short roots in its seedlings. Since GYY4137-mediated root stimulation was exclusively limited to *Rs* seedlings, but not in *Epi* and *Nr* mutants, shortening of *Rs* roots was not related to increased ethylene emission.

### The *Rs* mutation influences floral morphology

The major influence of the *Rs* mutation is on primary root development **(Yu, 1959;** https://tgrc-mvc.plantsciences.ucdavis.edu/Genes/Detail/Rs). The *Rs* phenotype at the later stage of plant development was not described. At the post-seedling stage, the shorter stature of *Rs* plants and altered leaf shape were similar to those reported for the *d* mutant in MicroTom (**Martı et al., 2006**). However, the *Rs* mutant is uniquely characterised by a compact inflorescence, increased inflorescence number, and abnormal floral morphology. The fruits originated from abnormal flowers had a twisted shape. As these phenotypes are absent in the other three mutants, they are likely specific to the *Rs* locus. Despite the altered fruit shape, the *Rs* mutation did not affect fruit carotenoid levels, which were nearly identical to those of the Chatham variety. In contrast, anthocyanin levels were elevated in *Rs* fruits, which may be due to the physiological stress induced by the mutation (**Georgiev, 1972**).

### Rs seedlings show Impaired Sulphur Metabolism

Since the H_2_S donors and L-cysteine stimulate the *Rs* root elongation, it can be assumed that *Rs* likely possess an altered metabolic homeostasis compared to the control plants. Since cysteine is the primary source of organic sulphur for all other S-compounds, a deficiency of it has wide effects, as multiple metabolic pathways are interlinked with S-containing compounds. Consistent with this notion, the metabolic profile of *Rs* seedlings was significantly different from that of AC. The metabolic alterations in *Rs* seedlings were closely similar to those reported in Arabidopsis subjected to sulphur starvation (**Nikiforova et al., 2005**). Similar to *Rs*, sulphur-starved Arabidopsis seedlings also exhibited significantly elevated levels of asparagine, β-alanine, GABA, glutamine, valine, putrescine, glucose, glucose 6-phosphate, and myoinositol 2-phosphate (**Dataset S8**). Analogous to Arabidopsis, a massive decrease in glycine levels was also observed in *Rs* seedlings.

In the cellular metabolome, sulphur is closely integrated with carbon/nitrogen metabolism. It is plausible that the reduction in sulphur-dependent amino acid synthesis redirects metabolic flux towards the branched-chain amino acid synthesis pathway. The increased levels of valine and leucine in *Rs* seedlings are consistent with this notion. In contrast to Arabidopsis, the magnitude of metabolite up- and down-regulation was several-fold higher in *Rs* (**Dataset S8**), indicating a metabolic block in *Rs* rather than an artificial block in Arabidopsis, such as the withholding of sulphur. Since the S-starvation of Soybean seedlings inhibits root elongation (**Zhou et al., 2024**), it can be construed that the metabolic block related to sulphur metabolism causes shortening of the root in *Rs*.

Ostensibly, the influence of *Rs* on root shortening is restricted to the seedling stage, as mature Rs plants show profuse roots compared to AC. Consistent with this, the *Rs* mature roots show fewer metabolic alterations compared to AC, with only upregulation of β-alanine and glucose overlapping with those of *Rs* seedlings. Moreover, metabolic alterations in the mature roots of Rs do not display the same severity as observed in the seedlings. It entails that either the initiation of the secondary roots obviates the S-starvation post-seedling stage, or the genetic lesion underlying root suppression is confined only to the seedling stage. Akin to *Rs* seedlings, seedlings of Arabidopsis double mutant *gso1;gso2* exhibit pronounced root growth defects, which can be rescued by sucrose supplementation (**Racolta et al., 2014**). The *gso1;gso2* mutant root phenotype is restricted to the seedling stage, and the plant shows normal root growth later on. Since the hydrolysis of seed reserves sustains early seedling growth, the genetic lesion in *Rs* may be either related to the transport of S-amino acids into the root or the assimilation of these amino acids in the root. Like *the gso1;gso2* mutant, the influence of the *Rs* mutation is temporally confined to the seedling stage, as evident by normal root development in later stages.

### Metabolic Alteration Subsides in Mature *Rs* Plants

After the seedling stage, upregulation of metabolites related to sulphur deficiency gradually subsided during the progression of plant growth. Accordingly, the 4^th^ node leaf of *Rs* showed upregulation of eight metabolites similar to S-starved Arabidopsis seedlings, while in the 8th node leaf, only two metabolites were similarly affected (**Nikiforova et al., 2005**). Despite the nodal position difference, an overlap in metabolic regulation was indicated as several metabolites were similarly up- or down-regulated in the above leaves. This implies that the metabolomic homeostasis of these leaves exhibits shared regulatory patterns (**Walker et al., 2023**). While the metabolic profiles of *Rs* leaves suggest a similarity to S-starved plants, it is plausible that other mutations, such as the *d* mutation, which influences leaf phenotype (**Martı et al., 2006**), may also contribute to metabolite alterations.

Despite showing a substantial number of abnormal flowers, the metabolome of *Rs* floral buds had the least differences from the control. The downregulation of the majority of metabolites in floral buds, including sugars, signifies increased energy demand for the growth of abnormal flowers. The decline of putrescine is reported during floral bud development; however, the massive downregulation in *Rs* is seemingly related to abnormal flower development (**Borghi et al., 2022**). It is plausible that abnormality in flower development arises due to improper S-assimilation. Comparison of nitrogen, phosphate, and sulphur deprivation in *Brassica rapa* and *Physalis philadelphica* revealed that reduction of flower size was exclusively due to S-deficiency (**Ausma et al., 2021**). The Microtom that harbours the *d* mutation, and also the *gs* and *h* mutants, does not display a flower phenotype. The abnormal flowers in *Rs* can also arise from misexpression of a homeotic gene such as WUS. The *Rs* fruit, with multiple locules, was somewhat similar to plants overexpressing the WUS gene, characterised by the absence of well-defined central septa and larger fruit size (**Li et al., 2017**). Therefore, it entails that the abnormal flower phenotype was either due to the *Rs* or in combination with other mutations.

Similar to floral buds, most metabolites in *Rs* fruits were downregulated, indicating that catabolism dominates the metabolic homeostasis. Considering that the reduction of fumarate and malate levels leads to a decrease in fruit size (**Centeno et al., 2011**), the upregulation of cis-aconitate, fumarate, and malate levels may be linked with the enlarged horizontal perimeter of *Rs* fruits. The downregulation of eight amino acids, three sugars, and other metabolites such as putrescine, signifies a massive metabolic shift in *Rs* fruits. The reduction in levels of the above metabolites, particularly sugars, putrescine, and increased malate levels, is associated with a lowering in the organoleptic qualities of fruits (**Osorio et al., 2020**).

### A root meristem-specific sulfotransferase(s) is likely responsible for the *Rs* phenotype

The similarities in the metabolic profiles of *Rs* and the sulfur-deprived plants hinted that a gene involved in sulfur metabolism underlies the *Rs* phenotype. Genetic mapping established that the mutation causing the *Rs* phenotype is localised on chromosome 4 (**Kerr, 1972**). Since sulfotransferases *Solyc04g028390* and *Solyc04g028380* are localised on chromosome 4 and are exclusively expressed in the root meristem, mutations in their promoter may affect gene expression, in turn disrupting root meristem functioning. Promoter mutations are known to affect gene expression, leading to altered phenotypes. In tomato, a single nucleotide mutation in the promoter of the *ACS2* gene reduces its *in vivo* expression, leading to delayed fruit ripening (**Sharma et al., 2021**). However, because both sulfotransferases are expressed exclusively in the root meristem niche, conventional RT-PCR is unsuitable for quantifying expression changes. Consequently, the precise molecular mechanism through which altered sulfotransferase expression influences the *Rs* phenotype remains unresolved.

Emerging reports have suggested that mutations in the promoters can result in a dominant developmental phenotype (**Liu et al., 2023; Yong et al., 2024**). In melon, three promoter mutations in the *pll* gene (*bm7* mutant) change constitutive expression to light/temperature-inducible expression, and transform round leaves present in WT to deeply palmately lobed leaves (**Yong et al., 2024).** In cabbage, a single base pair deletion in the promoter of male-sterile gene *Ms-cd1* confers a dominant phenotype by disrupting the binding of ethylene response factor ERF1L (**Han et al., 2023**). Likewise, the three promoter mutations in *Solyc04g028390* create both gain and loss of binding sites for several transcription factors, which may alter its expression. Taken together, the correlation mentioned above supports the likelihood that a sulfotransferase (*Solyc04g028390*) could be the candidate gene for the *Rs* phenotype.

Parallel to *Solyc04g028390*, the RGF-encoding gene *Solyc04g005240*, whose homologous mutants (**AT5G60810**) in Arabidopsis exhibit postembryonic root development defects, shows similar root-meristem-localised expression (**Matsuzaki et al., 2010)**. The co-localised expression of S*olyc04g005240* and *Solyc04g028390* hints at the linkage between the two, but is not supported by current literature. The Arabidopsis RGF1 (**AT5G60810**) is activated by sulfation at the tyrosine residue mediated by a tyrosine sulfotransferase (TPST) (**AT1G08030**). Mutations in Arabidopsis TPST (**AT1G08030**) cause defective root development by inhibiting sulfation of RGF1 (**Zhou et al., 2010**). The tomato homolog of Arabidopsis TPST, sulfotransferase Solyc11g069520, is localised on chromosome 11 of tomato and bears no mutations in the promoter and genic region. Unlike the root meristem-specific expression of *Solyc04g028390*, the TPSTs of 10(**Figures S8-9)**. The lack of homology with Solyc11g069520 suggests that Solyc04g028390 seemingly does not have tyrosine sulfotransferase activity. Interestingly, among all tomato RGFs and RGF-RKs, only *Solyc04g005240* shows root meristem-specific gene expression (**Figure S10**). The colocalised expression of three sulfotransferases (**Group VI**) and *Solyc04g005240* in root meristem may not be coincidental and, in probability, has a functional linkage.

While the role of tyrosine sulfotransferases is well established by several studies, emerging research is attempting to elucidate the roles of other sulfotransferases in plant metabolism. While these sulfotransferases are not involved in the sulfation of growth regulatory peptides, such as RGF/CLE/phytosulfokines, they reportedly *in vitro* catalyse the sulfation of a broad spectrum of biomacromolecules, including brassinosteroids, flavonoids, glucosinolates, jasmonates, etc (**Hirschmann et al., 2014**). Evidence for the *in vivo* sulfation of the above macromolecules by sulfotransferases remains equivocal. In tomatoes, the detection of a variety of sulfated brassinosteroids correlated with the occurrence of sulfotransferases; however, such a correlation was absent in Cucurbitaceae (**Han et al., 2025**). The likely association of *Solyc04g028390* expression with the root suppression phenotype suggests that the individual sulfotransferases may act in a tissue-specific fashion.

Notwithstanding the above, the association of *Solyc04g028390* with the *Rs* mutant phenotype is provisional, and more evidences are needed to support a firm linkage. The sulfotransferases require 3’-phosphoadenosine 5’-phosphosulfate (PAPS) for the sulfation of target molecules (**Hirschmann et al., 2014**). Neither H_2_S nor L-cysteine is a substrate or product in enzymatic reactions that generate PAPS in plants. Therefore, the observed rescue of root growth by H_2_S and L-cysteine is indirect, either by influencing the formation of PAPS or via some aspect related to H_2_S signalling. The evidence supports H_2_S signalling, as exogenous L-cysteine, which stimulated *Rs* root elongation, can generate H_2_S through its degradation by cysteine desulfurase (**Álvarez et al., 2010)**. Thus, it is likely that a sulfotransferase and H_2_S signalling, acting in concert or independently, may contribute to the *Rs* phenotype.

It is also plausible that promoter and genic mutations in adenylyl-sulfate kinase (*Solyc10g080680*) reduce the availability of PAPS, thus exacerbating the *Rs* phenotype. Considering that *Solyc10g080680* is expressed in all tissues of plants (**Figure 7**), the global influence of sulfur metabolism in different organs of *Rs* plants may arise from reduced supply of PAPS. Such an influence on sulfur metabolism may not be limited to adenylyl-sulfate kinase, as genome sequencing revealed a considerable number of mutations in *Rs* across the genome.

In all probability, the influence on sulfur metabolism of the *Rs* mutant and its morphological defects, particularly during fruit development, may also involve other genes that bear mutations in the gene and/or in the promoter region. The disruptions in sulphur assimilation cause a massive shift in the metabolite profile, as sulphur is tightly integrated with carbon/nitrogen metabolism and antioxidant systems. Our study reveals a likely relationship between the localised expression of sulfotransferases (*Solyc04g028380* and *Solyc04g028390*) in the root meristem and the functioning of the root meristem in tomato. The interconnecting process between the above sulfotransferases and the root meristem growth factor in tomato requires further detailed investigation.

## Materials and Methods

### Plant Materials and Chemicals

Tomato (*Solanum lycopersicum*) cv. Ailsa Craig (AC), obtained from Dr. Richard Kendrick (Wageningen University, The Netherlands), was used as the control. The *Rs* (LA1796; background genotype unknown), *Epi* (LA2089; background VFN8), and *Nr* (LA3537; background AC) mutants were sourced from the Tomato Genetics Resource Center (TGRC), UC Davis, California, USA (https://tgrc.ucdavis.edu/). The Chatham cultivar was provided by Dr. Maria Ivanchenko (Oregon State University, USA). Morpholin-4-ium 4-methoxy phenyl(mopholino) phosphinodithioate (GYY4137) was procured from Cayman chemicals (Cat.No.13345). Sodium hydrosulfide (NaHS, Cat. No.161527), DL-Propargylglycine (P7888), L-Cysteine (30089), and D-Cysteine (30095) were purchased from Sigma.\

### Germination and growth conditions

Seeds were surface-sterilised with 20% (v/v) sodium hypochlorite for 5–10 minutes, thoroughly rinsed with distilled water, and germinated in the dark at 25 ± 2°C on moist filter paper placed in plastic boxes (9.5 *l*× 9.5 *w*× 5 *h* cm). Upon radicle emergence, the sprouted seeds were used for experiments. They were transferred to vertically oriented 14 cm diameter (Ø) Petri plates lined with filter paper soaked with or without 12 mL of the respective chemical solution. The plates were then incubated either in darkness or under white light (100 µmol m⁻² s⁻¹) at 25 ± 2 °C. Seedling age was recorded from the time of radicle emergence. After five days, the lengths of the root and hypocotyl were measured.

For plant propagation, post-radicle emergence seeds were transferred to a vermiculite–peat mixture (Karnataka Explosives Ltd., Bangalore, India). After two weeks, seedlings were transplanted to red loam soil in pots and maintained in a greenhouse under natural day/night cycles. Comparisons between wild-type and mutant lines were performed using seed lots from the same harvest, grown together under identical conditions.

### Biochemical Characterization

For chlorophyll estimation, leaves were harvested from the 8th node of two-month-old plants. The chlorophyll content was estimated using the method described by **Porra et al. (1989).** The ethylene measurement from the 12-day-old light-grown seedlings was carried out as described in **Sharma et al. (2021).** Anthocyanins were extracted in acidified methanol (1% HCl) and quantitatively estimated by measuring the absorbance at 535 nm (**Harborne, 1967**). The carotenoid profiling was carried out following the procedure of **Gupta et al. (2015)**

### Metabolome Profiling

For metabolic profiling, unopened floral buds (10-day-old) were collected from the first inflorescence truss of the plant. The mature roots were collected by uprooting 3-month-old plants and carefully removing the surrounding soil. The leaves were collected from the 4th node (4–5-week-old plants) and the 8th node (8-9-week-old plants) of plants. The fruits for analysis were collected at the red-ripe stage of ripening from the first two trusses.

The primary metabolite extraction and analysis from seedlings, mature roots, leaves, floral buds, and fruits were carried out using a Leco Pegasus GC-MS, following a protocol modified from **Roessner et al. (2000),** as described in **Bodanapu et al. (2016).** Only metabolites with a ≥1.5-fold change (log2 ± 0.584) and p-value ≤0.05 were considered to be significantly different. Principal Component Analysis (PCA) was performed using Metaboanalyst 5.0 (http://www.metaboanalyst.ca/). Heat maps were generated using Morpheus (https://software.broadinstitute.org/morpheus/).

### Whole Genome Sequencing (WGS)

Whole-genome sequencing and bioinformatic analysis of the *Rs* mutant were performed according to the general workflow established in **Gupta et al. (2024)**. Since the genetic background of *Rs* is unknown (**Table S3**), the reads from whole-genome sequencing of *Rs* were aligned to the *S. lycopersicum* cv Heinz reference genome (SL3.0).

Sequencing was conducted by GeneWiz Inc. (NJ, USA) on an Illumina HiSeqX platform, yielding approximately 200 million paired-end (2 × 150 bp) reads (31.5 Gb), with over 29 Gb achieving a Q30 quality score. Raw reads were quality-filtered using **fastp (v0.19.5)** with the following parameters: -M 30 -3 -5. High-quality reads were then aligned to the Heinz SL3.0 genome using **BWA-MEM (v0.7.17)** (**Li and Durbin, 2009**). Variant calling and subsequent processing followed the methodology described in **Gupta et al. (2020).** Since the wild-type background of *Rs* is unknown (**Table S3**), the background variant subtraction was not performed. The resulting variants were annotated with the **SIFT4G** algorithm, using the SIFT4G-ITAG3.2 reference database (**Gupta et al., 2020**) to predict the impact of amino acid substitutions. Substitutions with SIFT scores ≤0.05 were classified as deleterious. Promoter analysis was performed using PCbase (http://pcbase.itps.ncku.edu.tw/) and loss or gain of transcription factor binding sites were retrieved (**Table S2**).

### Gene Expression Profiling, promoter analysis and Phylogeny Tree

For promoter analysis of *Solyc04g028380*, *Solyc04g028390*, and *Solyc04g064940*, a 2-kb region upstream of the ATG start codon was selected for each gene. Both wild-type (WT) and mutant promoter sequences were analyzed using PCbase 2.0 (**Chow et al 2024**) (https://pcbase.itps.ncku.edu.tw/promoter_analysis.php) with the *Arabidopsis thaliana* and *Solanum lycopersicum* databases to identify potential transcription factor binding sites. Additional transcription factor binding sites prediction was performed using PlantCARE (**Lescot et al., 2002**; http://bioinformatics.psb.ugent.be/webtools/plantcare/html/). To compare WT and mutant sequences and identify variations in TFBS regions, pairwise sequence alignment was performed using Multalin (http://multalin.toulouse.inra.fr/multalin/) (**Corpet 1988).**

The gene expression data for S-metabolism-related genes in various organs of tomato were obtained from the ConKet platform (https://conekt.sbs.ntu.edu.sg) as mean expression values (**Proost and Mutwil, 2018**). The relative expression values for each gene were normalized to their maximum expression level observed in any tissue (set to 1). The heatmaps were generated using relative expression values using Morpheus (https://software.broadinstitute.org/morpheus/). Amino acid sequences of sulfotransferase precursor peptides from Arabidopsis and tomato were first aligned with MAFFT software (https://mafft.cbrc.jp). This multiple sequence alignment was used as input for phylogenetic reconstruction with IQ-TREE software (http://iqtree.cibiv.univie.ac.at/) under the Maximum-likelihood criterion, with branch support assessed from 1,000 bootstrap replicates. The resulting tree was visualised as a circular phylogeny using the iTOL online tool (https://itol.embl.de).

## Supporting information

NaHS also elicited the hypocotyl elongation of the Rs seedlings, while NaHS-induced stimulations were lacking in AC

albeit at 0.1 mM and above concentrations of L-cysteine

GYY4137-treated Epi and Nr mutant seedlings did not elicit root elongation

Consistent with this, the anthocyanin level in Rs fruits was three times higher than in Chatham

Venn analysis indicated only a few overlapping up- or down-regulated metabolites between different developmental stages

nitrogen-rich compounds (glutamine, asparagine) indicated significant changes in amino acid metabolism

Solyc04g028390 protein within a unique clade of four sulfotransferases on tomato chromosome 4

tomato (Solyc11g069520) and Arabidopsis (AT1G08030) are expressed in almost all plant organs

tomato (Solyc11g069520) and Arabidopsis (AT1G08030) are expressed in almost all plant organs

and RGF-RKs, only Solyc04g005240 shows root meristem-specific gene expression

was the most promising candidate, as it contained three insertions and a transversion in its promoter region

binding sites of several transcription factors and, in parallel, added binding sites for additional TFs

the presence of three secondary mutations, which may also influence the observed phenotypes

The total carotenoid level and the distribution of carotenoids in Rs were nearly identical to those in Chatham

compared to AC, showing close clustering within Rs and clear separation from AC

only a few overlapping up- or down-regulated metabolites between different developmental stages

where 23 metabolites were upregulated and 21 were downregulated

with the Heinz reference genome identified mutations in several sulfur-related genes

several transcription factors and, in parallel, added binding sites for additional TFs

in the Rs genome on metabolism, we examined their expression in various organs of tomato.

GABA, glutamine, valine, putrescine, glucose, glucose 6-phosphate, and myoinositol 2-phosphate

## Acknowledgements

We thank Dr Maria Ivanchenko, Oregon State University, for providing us with the seeds of the Chatham cultivar.

## >Funding

This work was supported by the Department of Biotechnology (DBT), India grants, BT/PR11671/PBD/16/828/2008, BT/PR/7002/PBD/16/1009/2012, and BT/COE/34/SP15209/2015 to RS and YS and BT/INF/22/SP44787/2021 to YS and RS. We also acknowledge the funding from DBT-BUILDER (BT/INF/22/SP41176/2020) and MHRD-

IOE (RC5-22-019) to the laboratory. The Repository of Tomato Genomics Resources is a DBT-SAHAJ National facility.

## Supplementary Information

Figure S1: Influence of H_2_S donors and scavengers on the elongation of the root.

Figure S2: Influence of L-cysteine on *Rs* and Chatham root.

Figure S3: Influence of GYY(4137) and glutathione on *Nr, epi,* and *Rs* mutants of tomato.

Figure S4. Characterization of Chatham and *Rs* fruits.

Figure S5. Six-set Venn diagram representation of differentially expressed metabolites in different organs of *Rs*.

Figure S6. Metabolite profiles of different organs of the *Rs* mutant.

Figure S7. Phylogenetic relationship of sulfotransferase (SOT) proteins in *Solanum lycopersicum* and *Arabidopsis thaliana*.

Figure S8. Tissue-specific expression profile of sulfotransferase genes in tomato. Figure S9. Tissue-specific expression profile of sulfotransferase genes in Arabidopsis. Figure S10. Tissue-specific expression profile of RGF and RGF-RK genes in tomato.

Table S1. Mutations identified in the promoter regions of Solyc04g028380 and Solyc04g028390.

Table S2. Loss or gain of transcription factor binding sites due to promoter mutations.

Table S3. The accession and mutant gene details of LA1796.

Dataset S1. The carotenoid levels in red-ripe fruits of Chatham and *Rs*.

Dataset S2. List of metabolites identified at different stages of *Rs* and control plants.

Dataset S3. A six-set Venn analysis showing down-regulated metabolites between different developmental stages

Dataset S4. Significantly up- and downregulated metabolites at one or more stages in *Rs* plants relative to respective controls.

Dataset S5. List of mutated tomato genes related to sulfur metabolism and transport.

Dataset S6. Loss/gain in transcription factor binding site due to promoter mutations.

Dataset S7. Expression of sulfur-related genes in different organs/tissues of tomato and Arabidopsis

Dataset S8. Comparison of metabolite levels in *Rs* seedlings with Sulfur-starved Arabidopsis seedlings.

