## Supplementary material for "Root-Suppressed Phenotype of Tomato *Rs* Mutant is Seemingly Related to Expression of Root-Meristem-Specific Sulfotransferases": NaHS also elicited the hypocotyl elongation of the Rs seedlings, while NaHS-induced stimulations were lacking in AC

### Slide 1
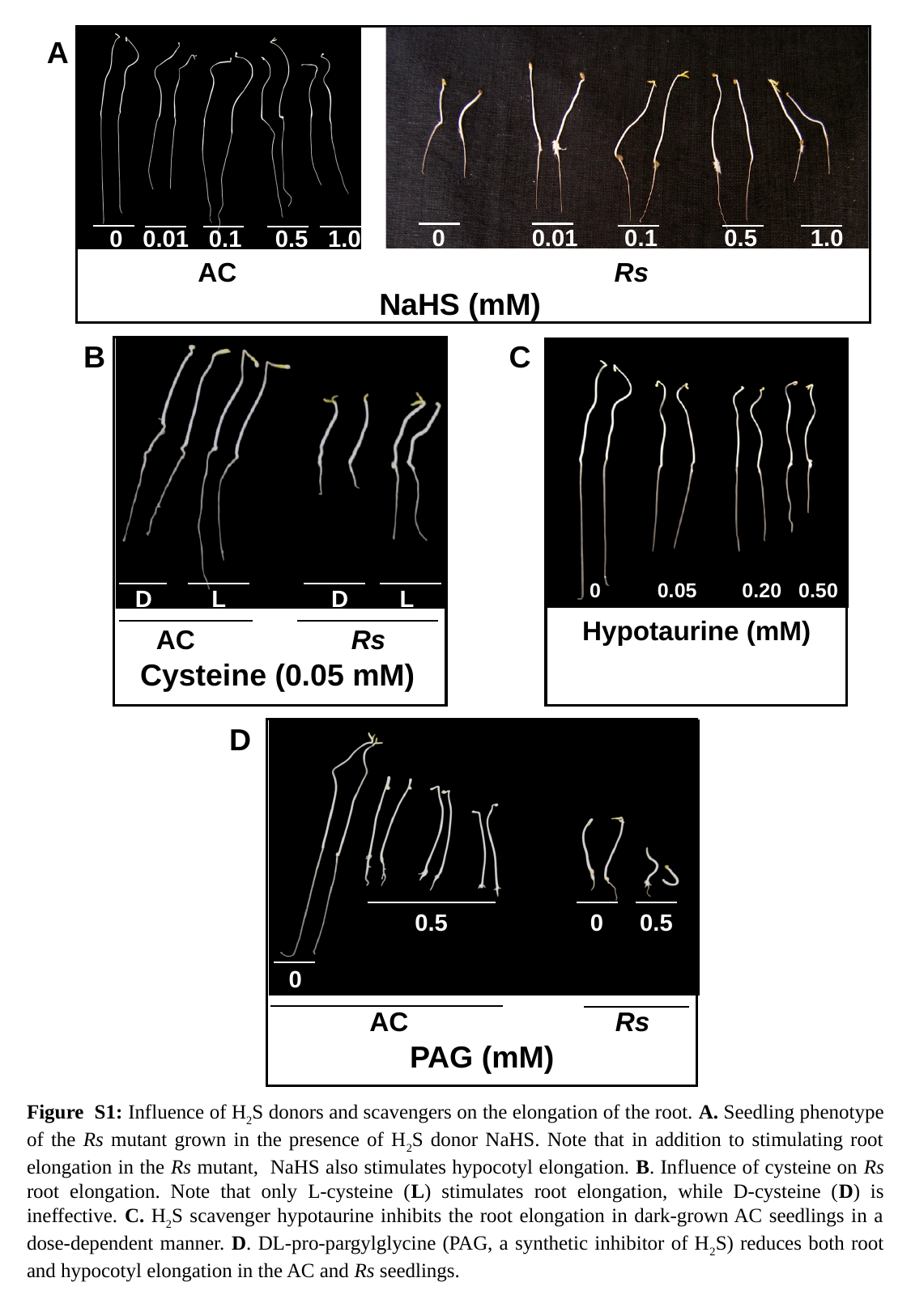

0 0.01 0.1 0.5 1.0
AC
0 0.01 0.1 0.5 1.0
Rs
NaHS (mM)
A
B
C
 D
 L
 D
 L
AC
Rs
Cysteine (0.05 mM)
 0 0.05 0.20 0.50
Hypotaurine (mM)
D
 0.5
 0
 0.5
 0
AC
Rs
PAG (mM)
Figure S1: Influence of H_2_S donors and scavengers on the elongation of the root. A. Seedling phenotype of the Rs mutant grown in the presence of H_2_S donor NaHS. Note that in addition to stimulating root elongation in the Rs mutant, NaHS also stimulates hypocotyl elongation. B. Influence of cysteine on Rs root elongation. Note that only L-cysteine (L) stimulates root elongation, while D-cysteine (D) is ineffective. C. H_2_S scavenger hypotaurine inhibits the root elongation in dark-grown AC seedlings in a dose-dependent manner. D. DL-pro-pargylglycine (PAG, a synthetic inhibitor of H_2_S) reduces both root and hypocotyl elongation in the AC and Rs seedlings.
