## Supplementary material for "Root-Suppressed Phenotype of Tomato *Rs* Mutant is Seemingly Related to Expression of Root-Meristem-Specific Sulfotransferases": albeit at 0.1 mM and above concentrations of L-cysteine

### Slide 1
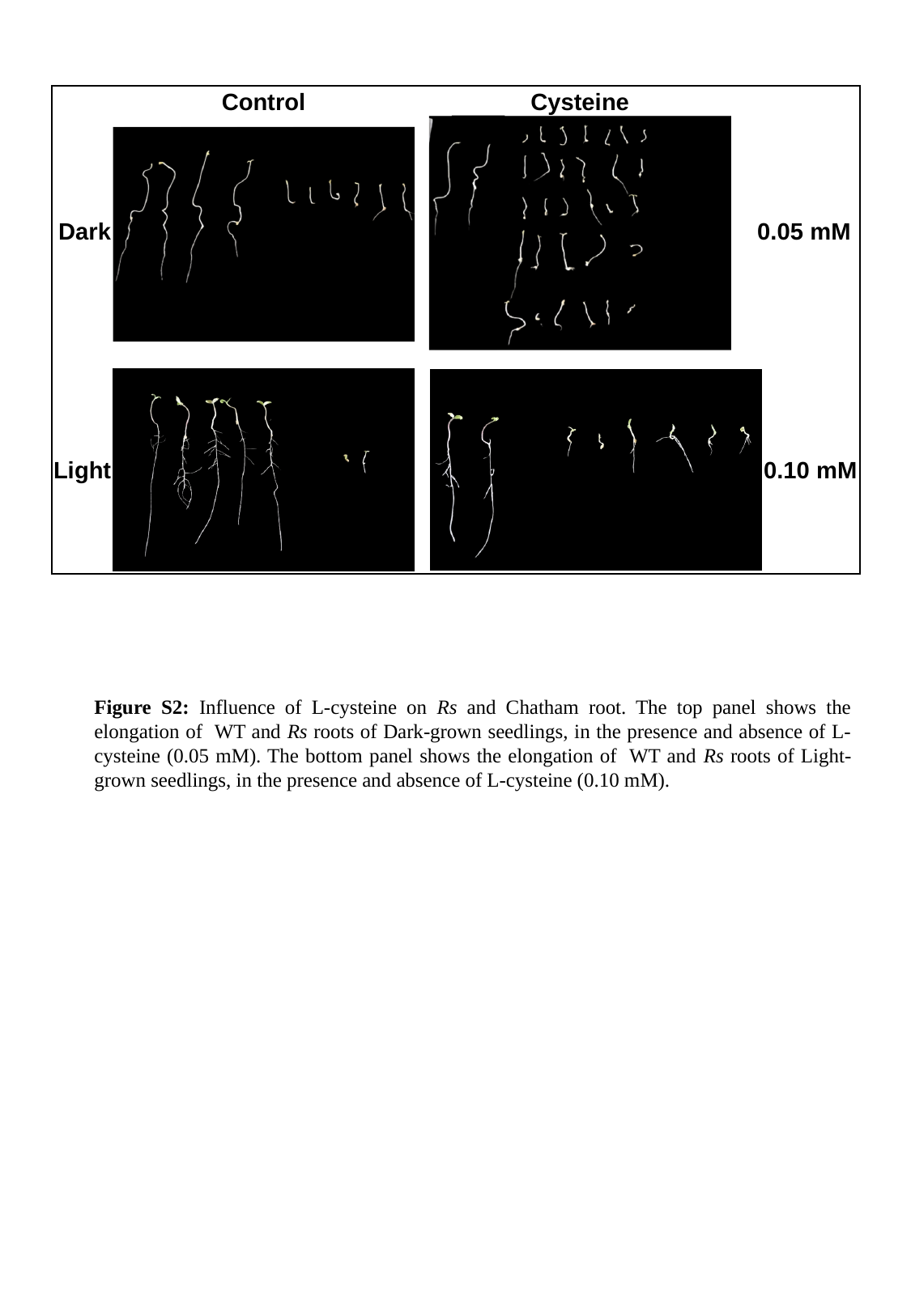

Control
Cysteine
Dark
0.05 mM
Light
0.10 mM
Figure S2: Influence of L-cysteine on Rs and Chatham root. The top panel shows the elongation of WT and Rs roots of Dark-grown seedlings, in the presence and absence of L-cysteine (0.05 mM). The bottom panel shows the elongation of WT and Rs roots of Light-grown seedlings, in the presence and absence of L-cysteine (0.10 mM).
