## Supplementary material for "Root-Suppressed Phenotype of Tomato *Rs* Mutant is Seemingly Related to Expression of Root-Meristem-Specific Sulfotransferases": GYY4137-treated Epi and Nr mutant seedlings did not elicit root elongation

### Slide 1
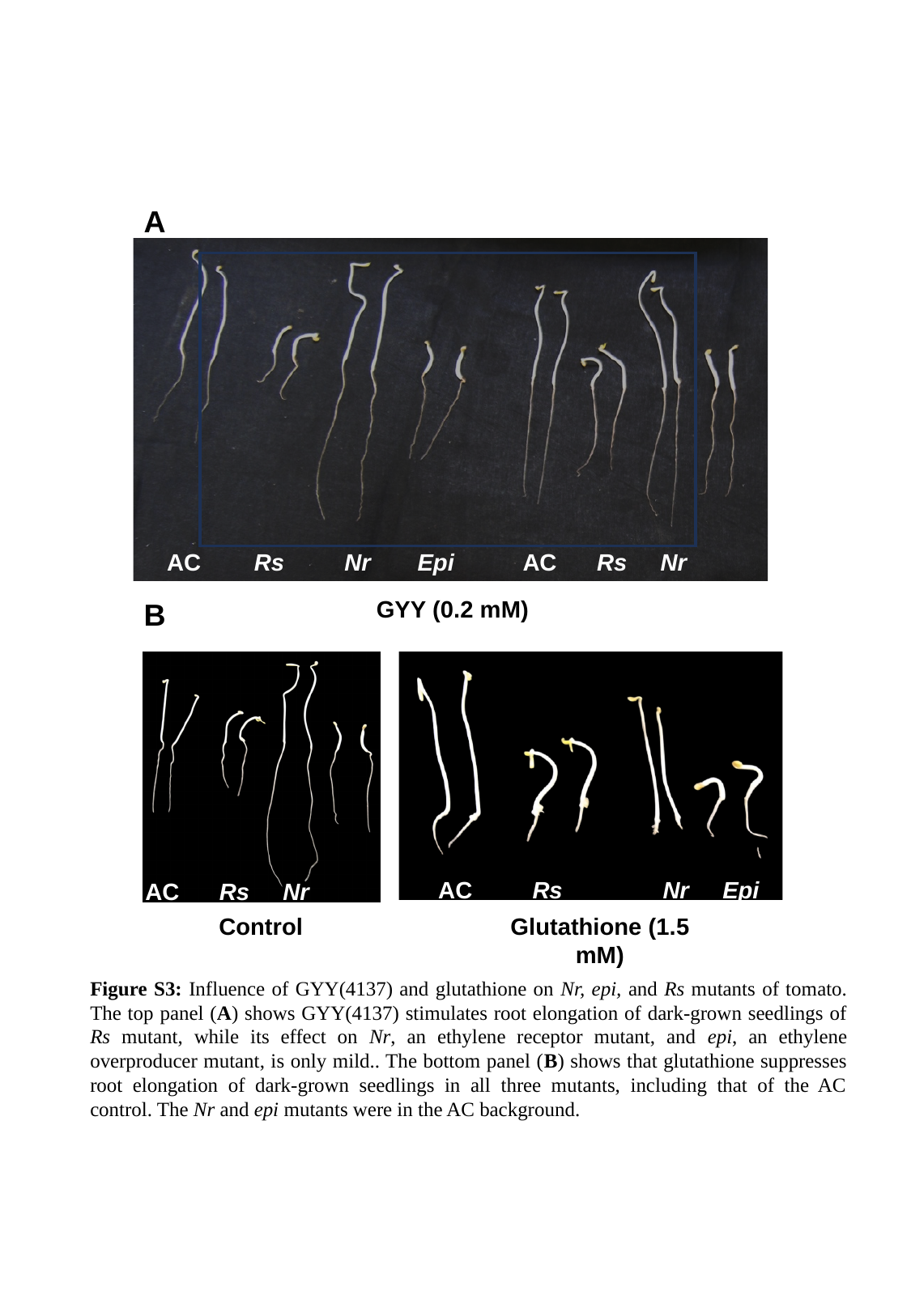

A
GYY (0.2 mM)
AC Rs Nr Epi
AC Rs Nr Epi
AC Rs Nr Epi
AC Rs Nr Epi
Control
Glutathione (1.5 mM)
B
Figure S3: Influence of GYY(4137) and glutathione on Nr, epi, and Rs mutants of tomato. The top panel (A) shows GYY(4137) stimulates root elongation of dark-grown seedlings of Rs mutant, while its effect on Nr, an ethylene receptor mutant, and epi, an ethylene overproducer mutant, is only mild.. The bottom panel (B) shows that glutathione suppresses root elongation of dark-grown seedlings in all three mutants, including that of the AC control. The Nr and epi mutants were in the AC background.
