## Supplementary material for "Root-Suppressed Phenotype of Tomato *Rs* Mutant is Seemingly Related to Expression of Root-Meristem-Specific Sulfotransferases": Consistent with this, the anthocyanin level in Rs fruits was three times higher than in Chatham

### Slide 1
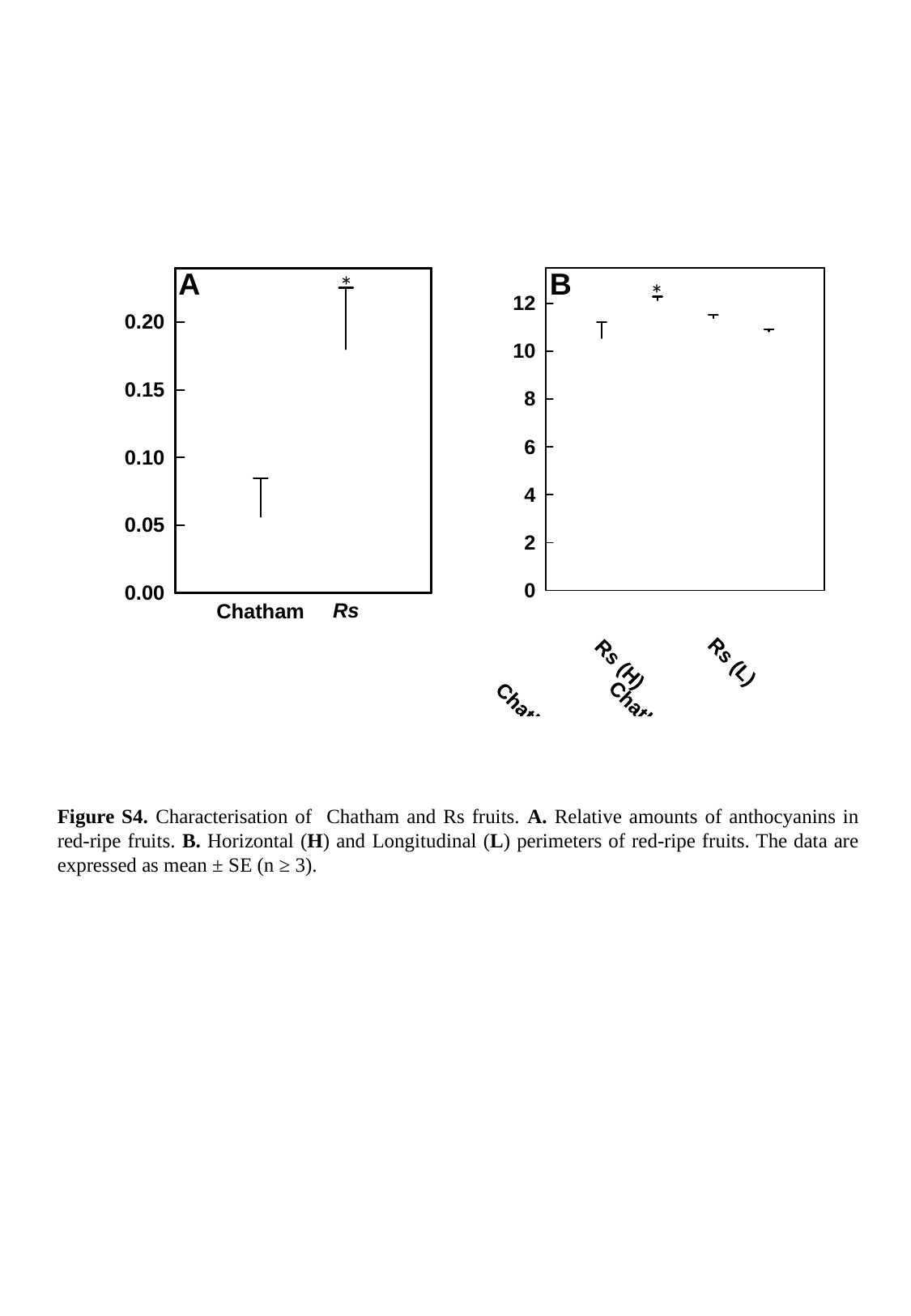

A
B
*
*
Figure S4. Characterisation of Chatham and Rs fruits. A. Relative amounts of anthocyanins in red-ripe fruits. B. Horizontal (H) and Longitudinal (L) perimeters of red-ripe fruits. The data are expressed as mean ± SE (n ≥ 3).
