## Supplementary material for "Root-Suppressed Phenotype of Tomato *Rs* Mutant is Seemingly Related to Expression of Root-Meristem-Specific Sulfotransferases": Venn analysis indicated only a few overlapping up- or down-regulated metabolites between different developmental stages

### Slide 1
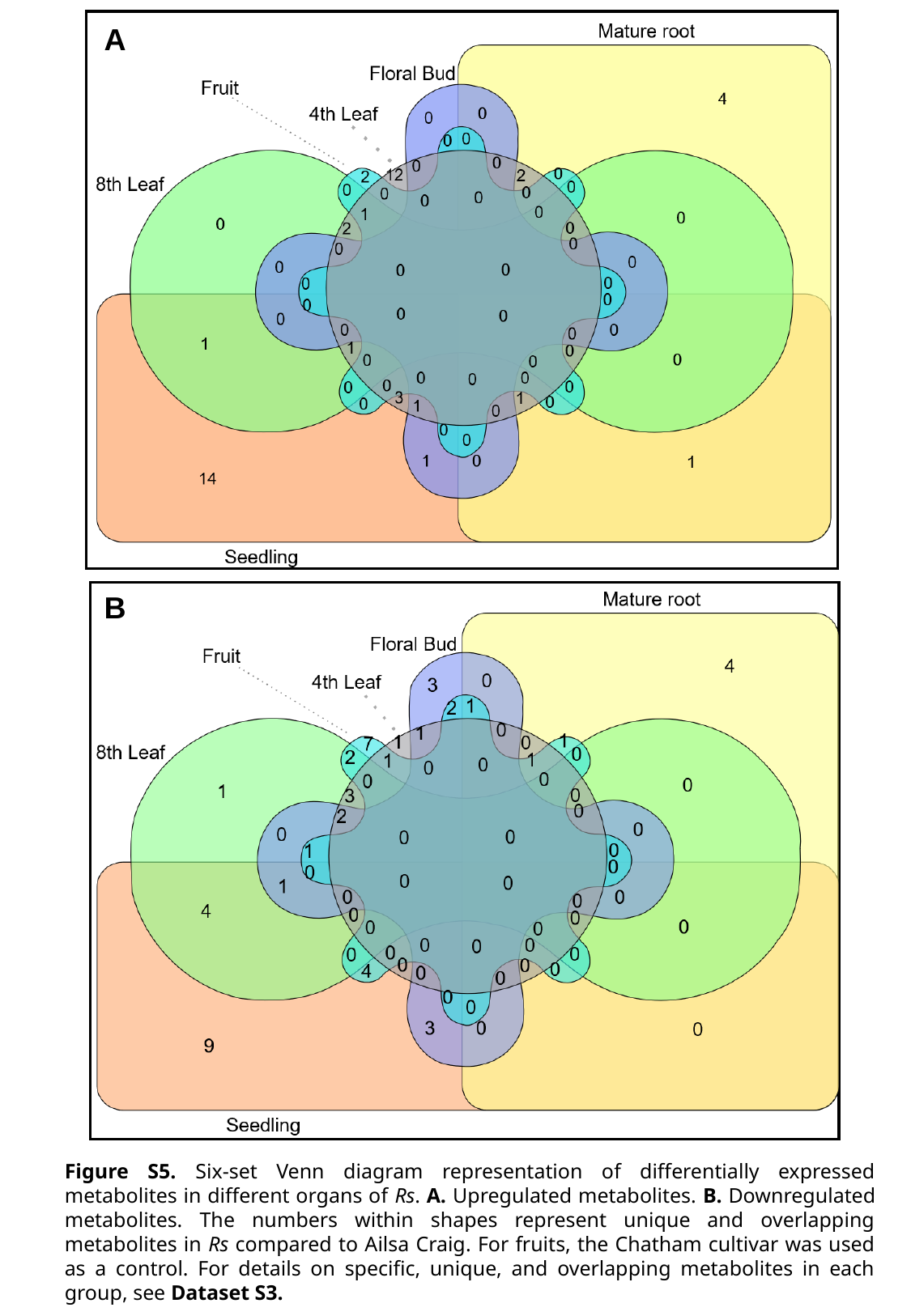

A
B
Figure S5. Six-set Venn diagram representation of differentially expressed metabolites in different organs of Rs. A. Upregulated metabolites. B. Downregulated metabolites. The numbers within shapes represent unique and overlapping metabolites in Rs compared to Ailsa Craig. For fruits, the Chatham cultivar was used as a control. For details on specific, unique, and overlapping metabolites in each group, see Dataset S3.
