## Supplementary material for "Root-Suppressed Phenotype of Tomato *Rs* Mutant is Seemingly Related to Expression of Root-Meristem-Specific Sulfotransferases": nitrogen-rich compounds (glutamine, asparagine) indicated significant changes in amino acid metabolism

### Slide 1
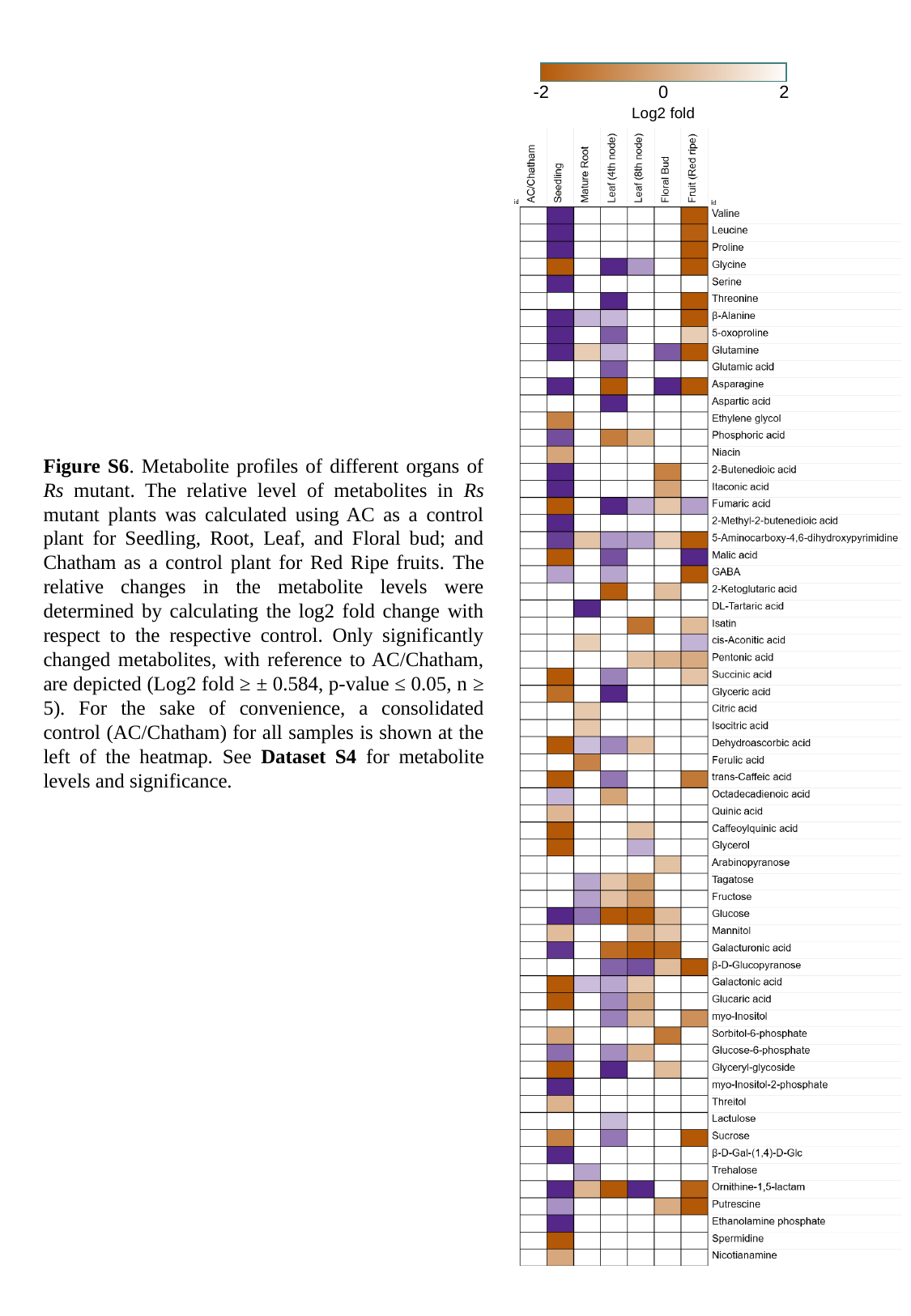

-2
0
2
Log2 fold
Figure S6. Metabolite profiles of different organs of Rs mutant. The relative level of metabolites in Rs mutant plants was calculated using AC as a control plant for Seedling, Root, Leaf, and Floral bud; and Chatham as a control plant for Red Ripe fruits. The relative changes in the metabolite levels were determined by calculating the log2 fold change with respect to the respective control. Only significantly changed metabolites, with reference to AC/Chatham, are depicted (Log2 fold ≥ ± 0.584, p-value ≤ 0.05, n ≥ 5). For the sake of convenience, a consolidated control (AC/Chatham) for all samples is shown at the left of the heatmap. See Dataset S4 for metabolite levels and significance.
