## Supplementary material for "Root-Suppressed Phenotype of Tomato *Rs* Mutant is Seemingly Related to Expression of Root-Meristem-Specific Sulfotransferases": Solyc04g028390 protein within a unique clade of four sulfotransferases on tomato chromosome 4

### Slide 1
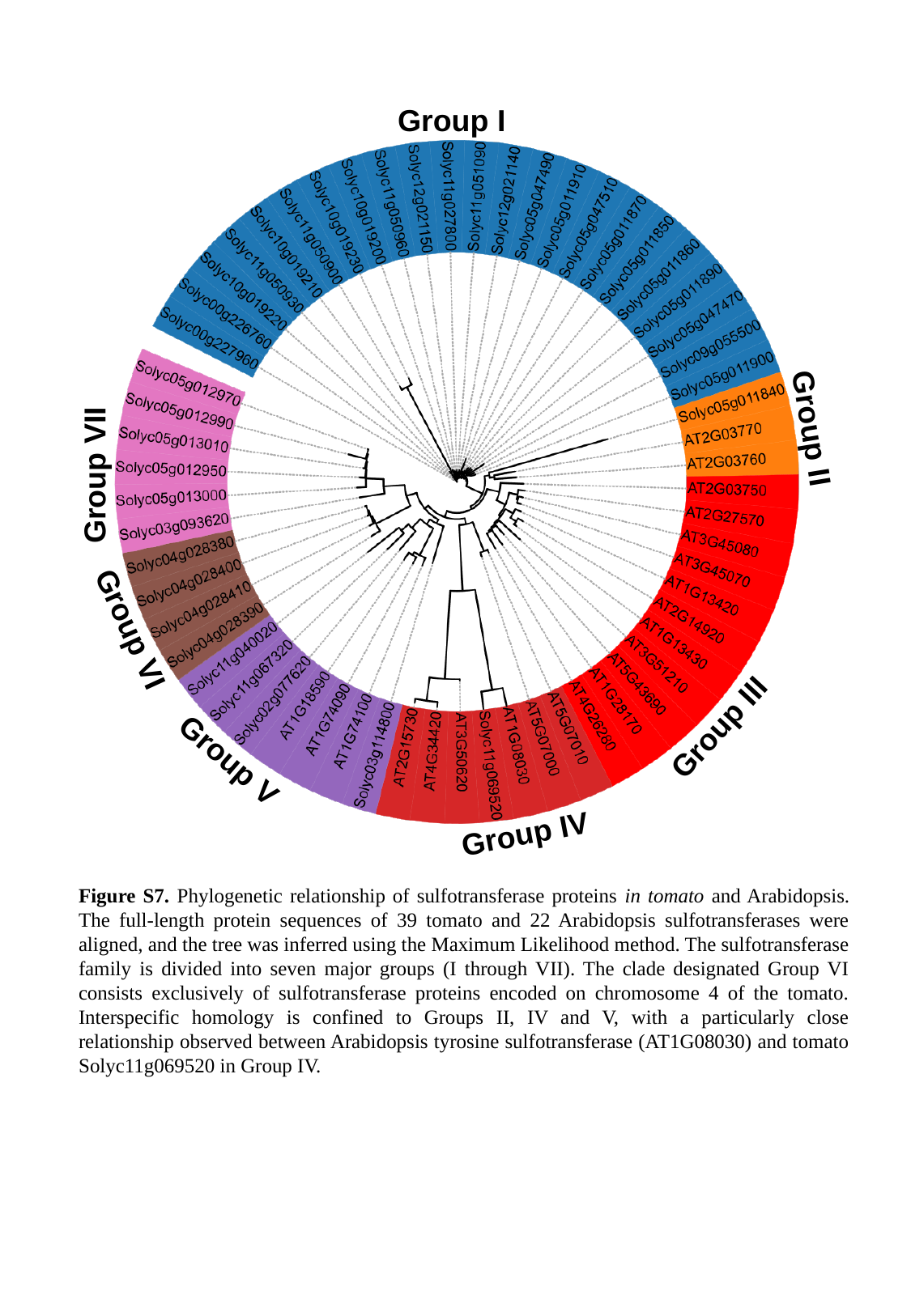

Group I
Group II
Group VII
Group VI
Group III
Group V
Group IV
Figure S7. Phylogenetic relationship of sulfotransferase proteins in tomato and Arabidopsis. The full-length protein sequences of 39 tomato and 22 Arabidopsis sulfotransferases were aligned, and the tree was inferred using the Maximum Likelihood method. The sulfotransferase family is divided into seven major groups (I through VII). The clade designated Group VI consists exclusively of sulfotransferase proteins encoded on chromosome 4 of the tomato. Interspecific homology is confined to Groups II, IV and V, with a particularly close relationship observed between Arabidopsis tyrosine sulfotransferase (AT1G08030) and tomato Solyc11g069520 in Group IV.
