## Supplementary material for "Root-Suppressed Phenotype of Tomato *Rs* Mutant is Seemingly Related to Expression of Root-Meristem-Specific Sulfotransferases": tomato (Solyc11g069520) and Arabidopsis (AT1G08030) are expressed in almost all plant organs

### Slide 1
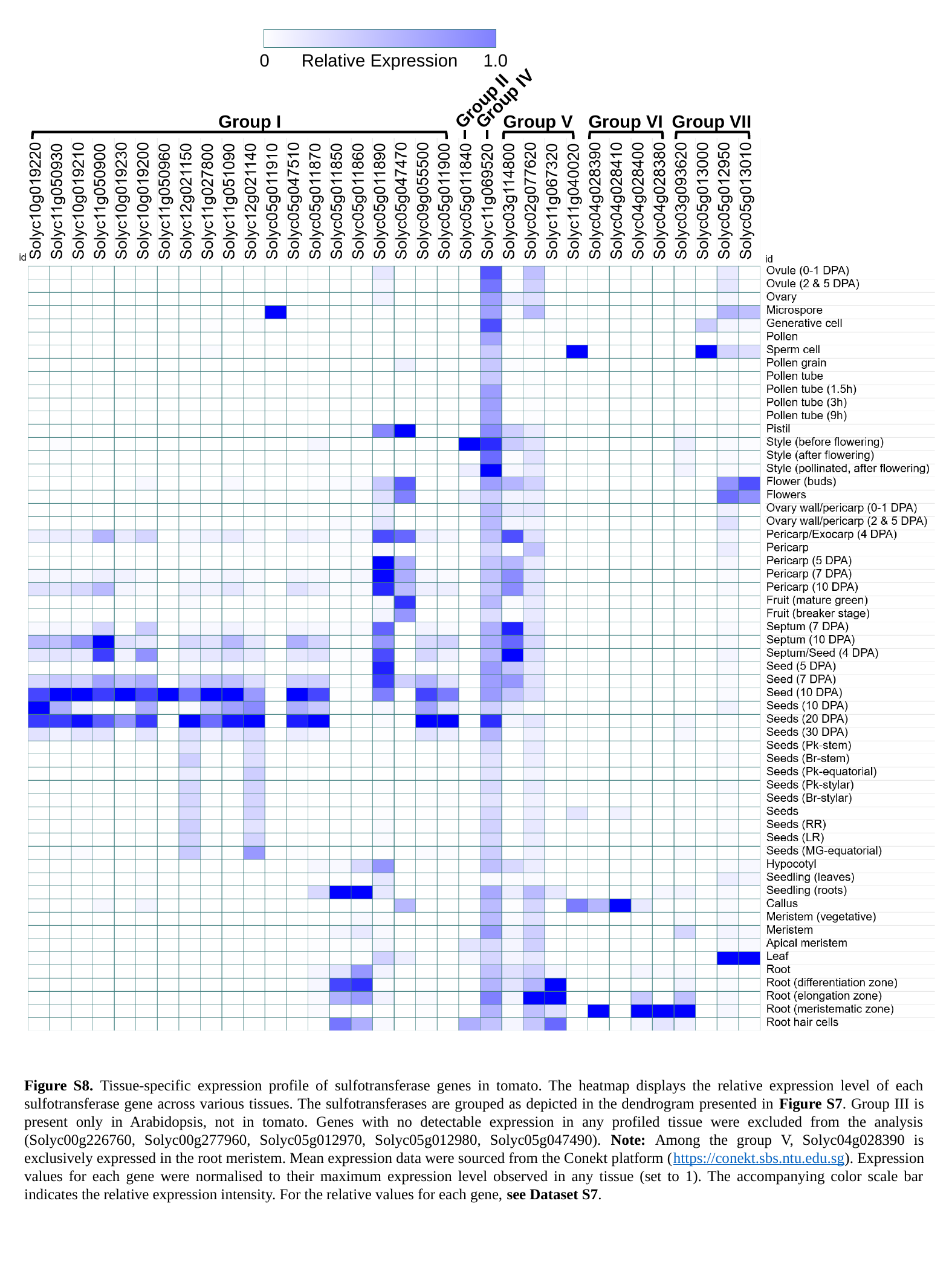

0
Relative Expression
1.0
Group II
Group IV
Group I
Group V
Group VI
Group VII
Figure S8. Tissue-specific expression profile of sulfotransferase genes in tomato. The heatmap displays the relative expression level of each sulfotransferase gene across various tissues. The sulfotransferases are grouped as depicted in the dendrogram presented in Figure S7. Group III is present only in Arabidopsis, not in tomato. Genes with no detectable expression in any profiled tissue were excluded from the analysis (Solyc00g226760, Solyc00g277960, Solyc05g012970, Solyc05g012980, Solyc05g047490). Note: Among the group V, Solyc04g028390 is exclusively expressed in the root meristem. Mean expression data were sourced from the Conekt platform (https://conekt.sbs.ntu.edu.sg). Expression values for each gene were normalised to their maximum expression level observed in any tissue (set to 1). The accompanying color scale bar indicates the relative expression intensity. For the relative values for each gene, see Dataset S7.
