## Supplementary material for "Root-Suppressed Phenotype of Tomato *Rs* Mutant is Seemingly Related to Expression of Root-Meristem-Specific Sulfotransferases": tomato (Solyc11g069520) and Arabidopsis (AT1G08030) are expressed in almost all plant organs

### Slide 1
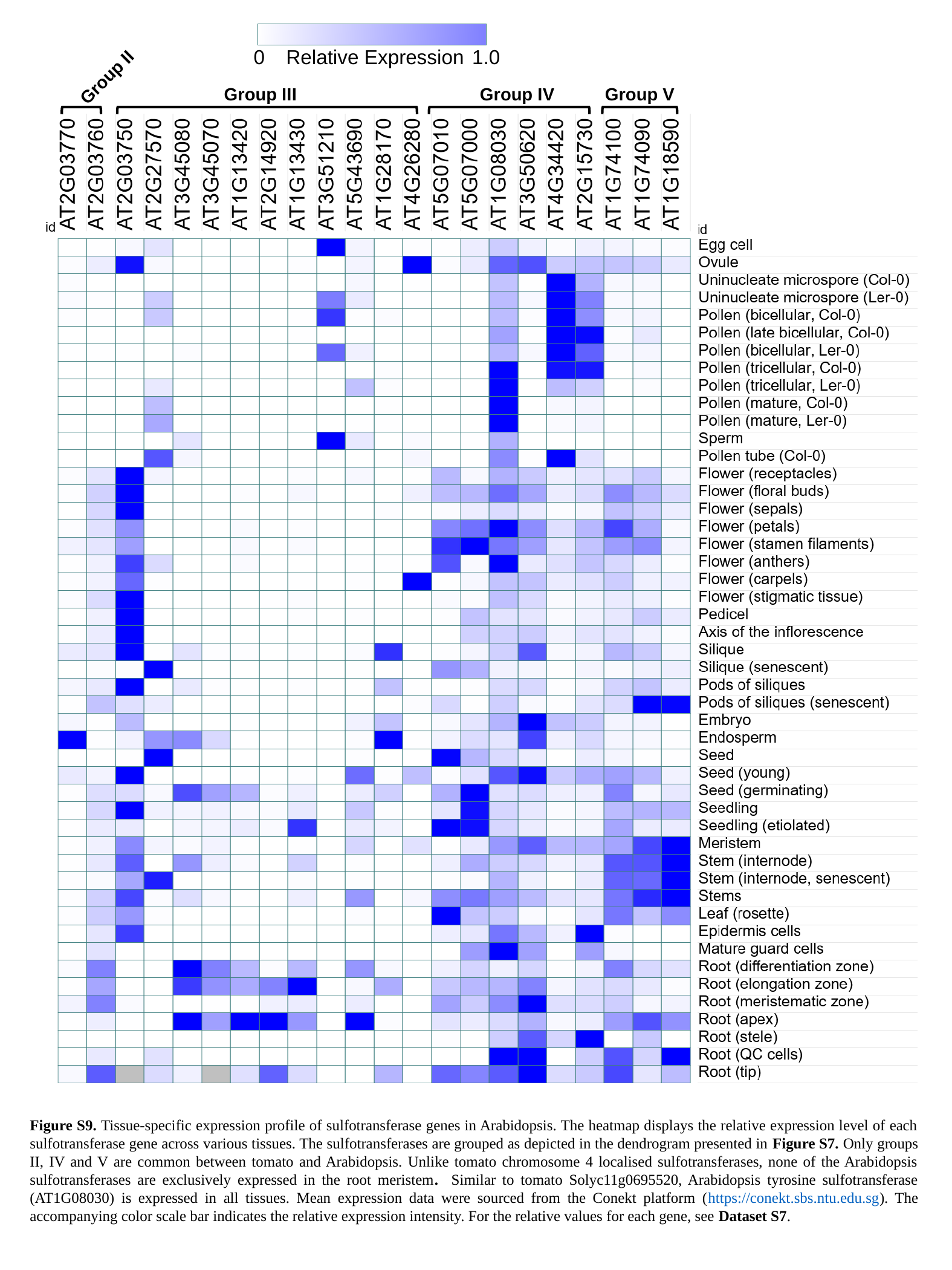

0
Relative Expression
1.0
Group II
Group III
Group IV
Group V
Figure S9. Tissue-specific expression profile of sulfotransferase genes in Arabidopsis. The heatmap displays the relative expression level of each sulfotransferase gene across various tissues. The sulfotransferases are grouped as depicted in the dendrogram presented in Figure S7. Only groups II, IV and V are common between tomato and Arabidopsis. Unlike tomato chromosome 4 localised sulfotransferases, none of the Arabidopsis sulfotransferases are exclusively expressed in the root meristem. Similar to tomato Solyc11g0695520, Arabidopsis tyrosine sulfotransferase (AT1G08030) is expressed in all tissues. Mean expression data were sourced from the Conekt platform (https://conekt.sbs.ntu.edu.sg). The accompanying color scale bar indicates the relative expression intensity. For the relative values for each gene, see Dataset S7.
