## Supplementary material for "Root-Suppressed Phenotype of Tomato *Rs* Mutant is Seemingly Related to Expression of Root-Meristem-Specific Sulfotransferases": and RGF-RKs, only Solyc04g005240 shows root meristem-specific gene expression

### Slide 1
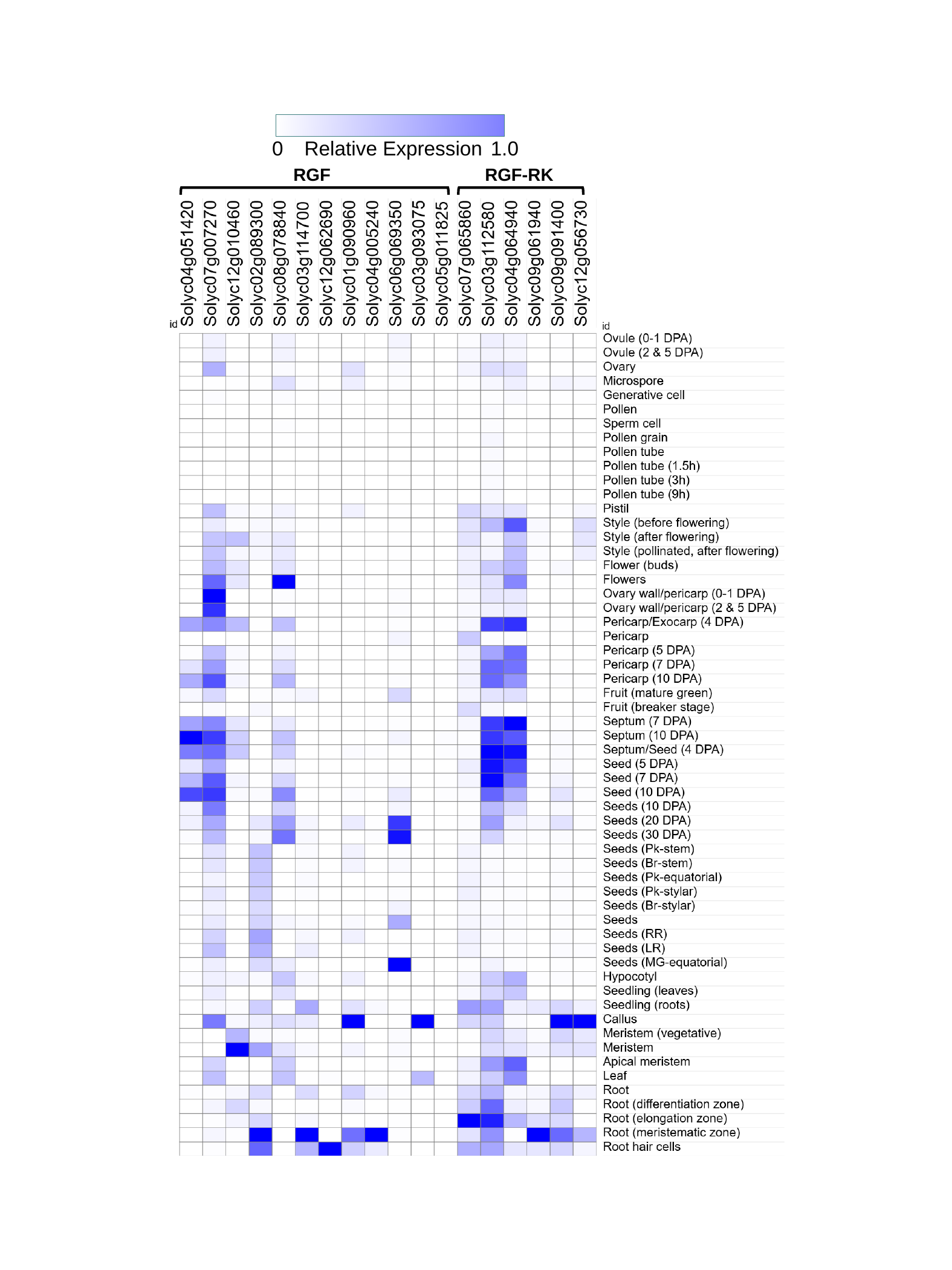

0
Relative Expression
1.0
RGF
RGF-RK

### Slide 2
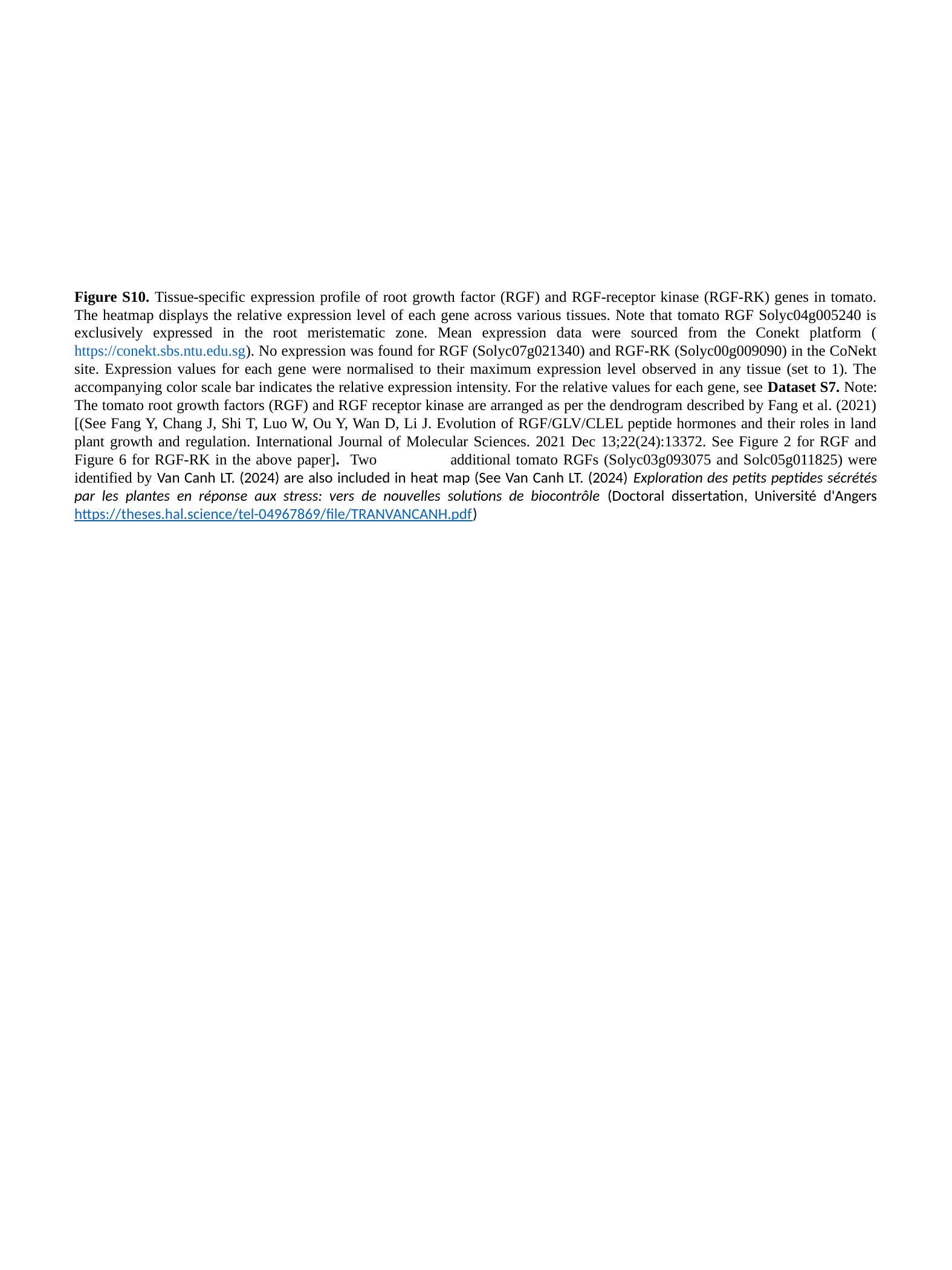

Figure S10. Tissue-specific expression profile of root growth factor (RGF) and RGF-receptor kinase (RGF-RK) genes in tomato. The heatmap displays the relative expression level of each gene across various tissues. Note that tomato RGF Solyc04g005240 is exclusively expressed in the root meristematic zone. Mean expression data were sourced from the Conekt platform (https://conekt.sbs.ntu.edu.sg). No expression was found for RGF (Solyc07g021340) and RGF-RK (Solyc00g009090) in the CoNekt site. Expression values for each gene were normalised to their maximum expression level observed in any tissue (set to 1). The accompanying color scale bar indicates the relative expression intensity. For the relative values for each gene, see Dataset S7. Note: The tomato root growth factors (RGF) and RGF receptor kinase are arranged as per the dendrogram described by Fang et al. (2021) [(See Fang Y, Chang J, Shi T, Luo W, Ou Y, Wan D, Li J. Evolution of RGF/GLV/CLEL peptide hormones and their roles in land plant growth and regulation. International Journal of Molecular Sciences. 2021 Dec 13;22(24):13372. See Figure 2 for RGF and Figure 6 for RGF-RK in the above paper]. Two	additional tomato RGFs (Solyc03g093075 and Solc05g011825) were identified by Van Canh LT. (2024) are also included in heat map (See Van Canh LT. (2024) Exploration des petits peptides sécrétés par les plantes en réponse aux stress: vers de nouvelles solutions de biocontrôle (Doctoral dissertation, Université d'Angers https://theses.hal.science/tel-04967869/file/TRANVANCANH.pdf)
