## Supplementary material for "Root-Suppressed Phenotype of Tomato *Rs* Mutant is Seemingly Related to Expression of Root-Meristem-Specific Sulfotransferases": was the most promising candidate, as it contained three insertions and a transversion in its promoter region

**Table S1.**  Mutations identified in the promoter regions of Solyc04g028380, Solyc04g028390 and Solyc04g064940. Bold, enlarged nucleotides in the mutant sequence indicate substitutions. Insertions in the mutant are represented by a hyphen (-) in the corresponding wild-type sequence. The complete promoter sequence alignment is shown on the following pages.

| **Mutants identified** | **Position of mutation** | **Change in Amino acid** | **Promoter element modified** |
| --- | --- | --- | --- |
| *Sulfotransferase*  (Solyc04g028380) | **A-393T** | Promoter | TGTTTA**A**AAATATA  **WT**  TGTTTA**T**AAATATA  ***mutant*** |
| *Sulfotransferase*  (Solyc04g028390) | **-530**  **T insertion** | Promoter | AATTTTTT**‑**AAACATAA  **WT**  AATTTTTT**T**AAACATAA  ***mutant*** |
|  | **A-1401T** | Promoter | TCCTTTTT**A**ATTTTT  **WT**  TCCTTTTT**T**AATTTTTT  ***mutant*** |
|  | **-1403,**  **-1404**  **AT insertion** | Promoter | TTTTTAA**‑‑**TTTTT  **WT**  TTTTTTA**AT**TTTTT  ***mutant*** |
|  | **A insertion**  **-1576** | Promoter | AAAAA**‑**CTAG  **WT**  AAAAA**A**CTAG  ***mutant*** |
| *Receptor protein kinase*  (Solyc04g064940) | **-1540,**  **-1541**  **TT insertion** | Promoter | TTTTTT--AAATTTT  **WT**  TTTTTT**TT**AAATTTT  ***mutant*** |

Sequence alignment of the promoter region of (a) *Solyc04g028380,* (b) *Solyc04g028390* and (c) *Solyc04g064940.* The ~2-kb promoter region of the wild-type (WT) was retrieved from the ITAG3.0 database and aligned with the *Rs* mutant sequence using Multalin (Corpet, 1988). Nucleotide differences between the WT and mutant sequences are highlighted in blue and black. The specific mutated region is shown on the right side of the promoter sequence.

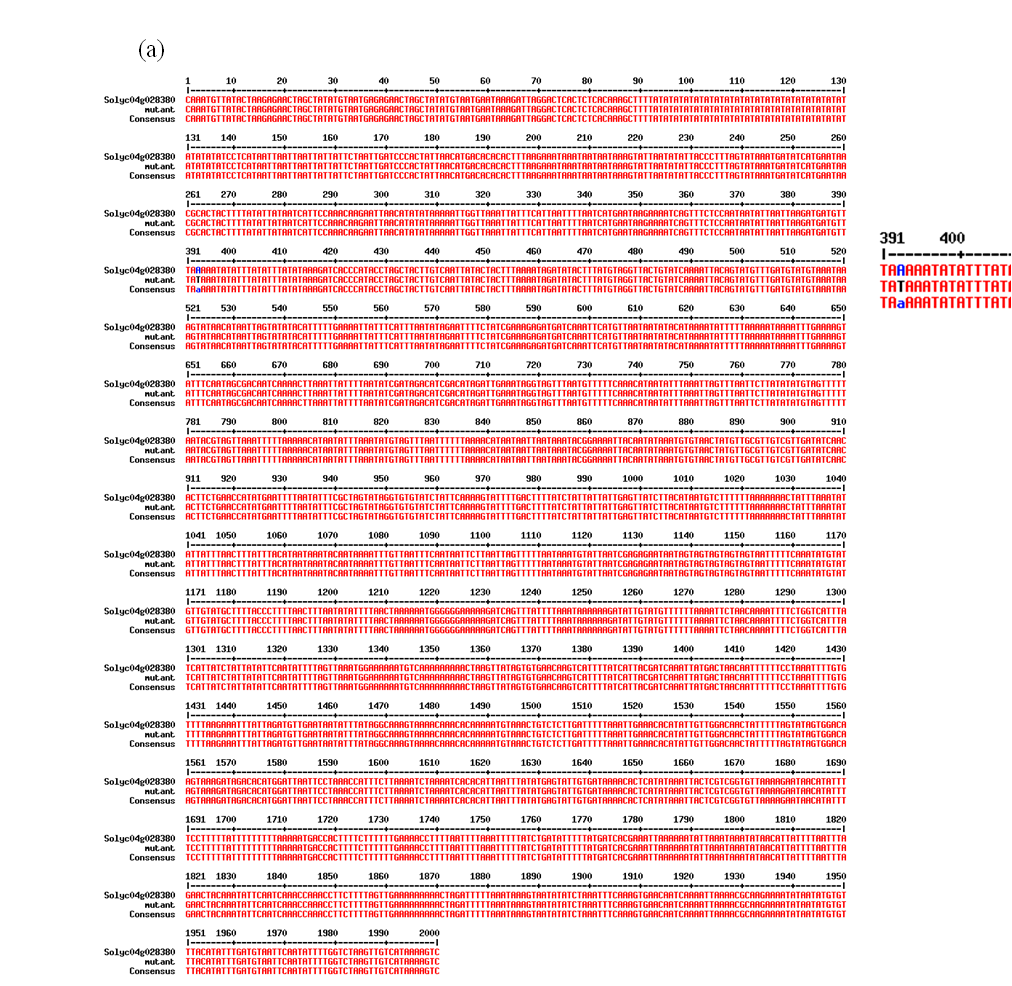

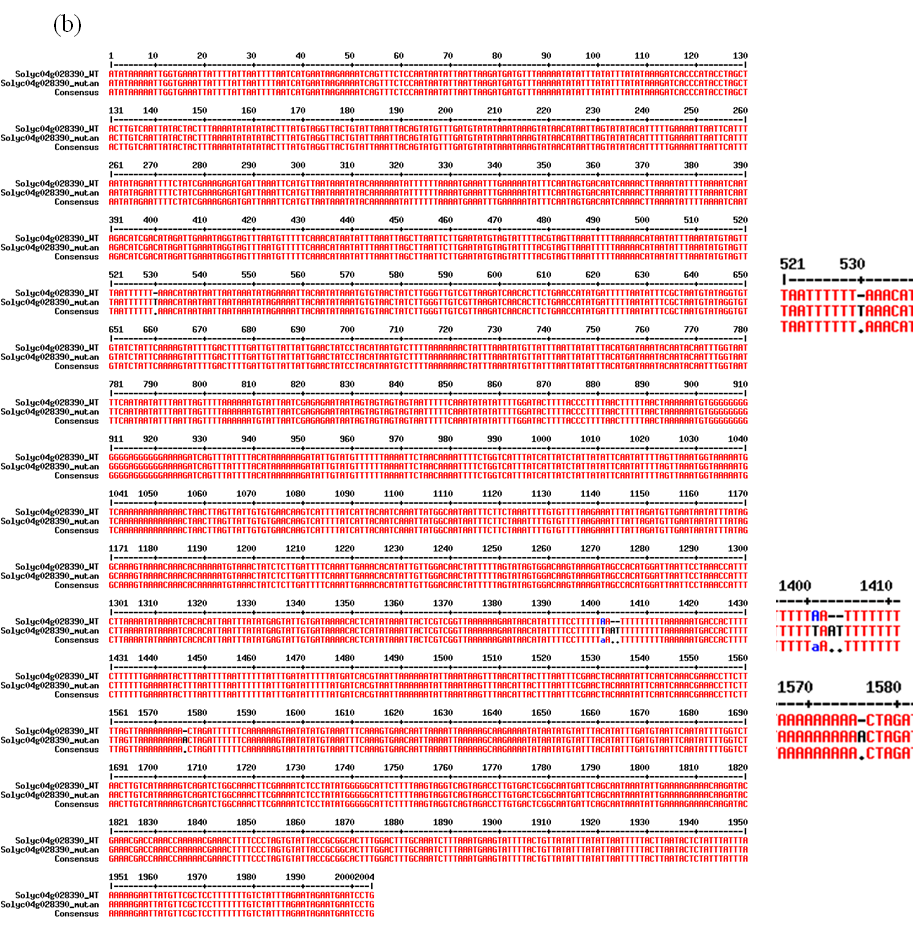

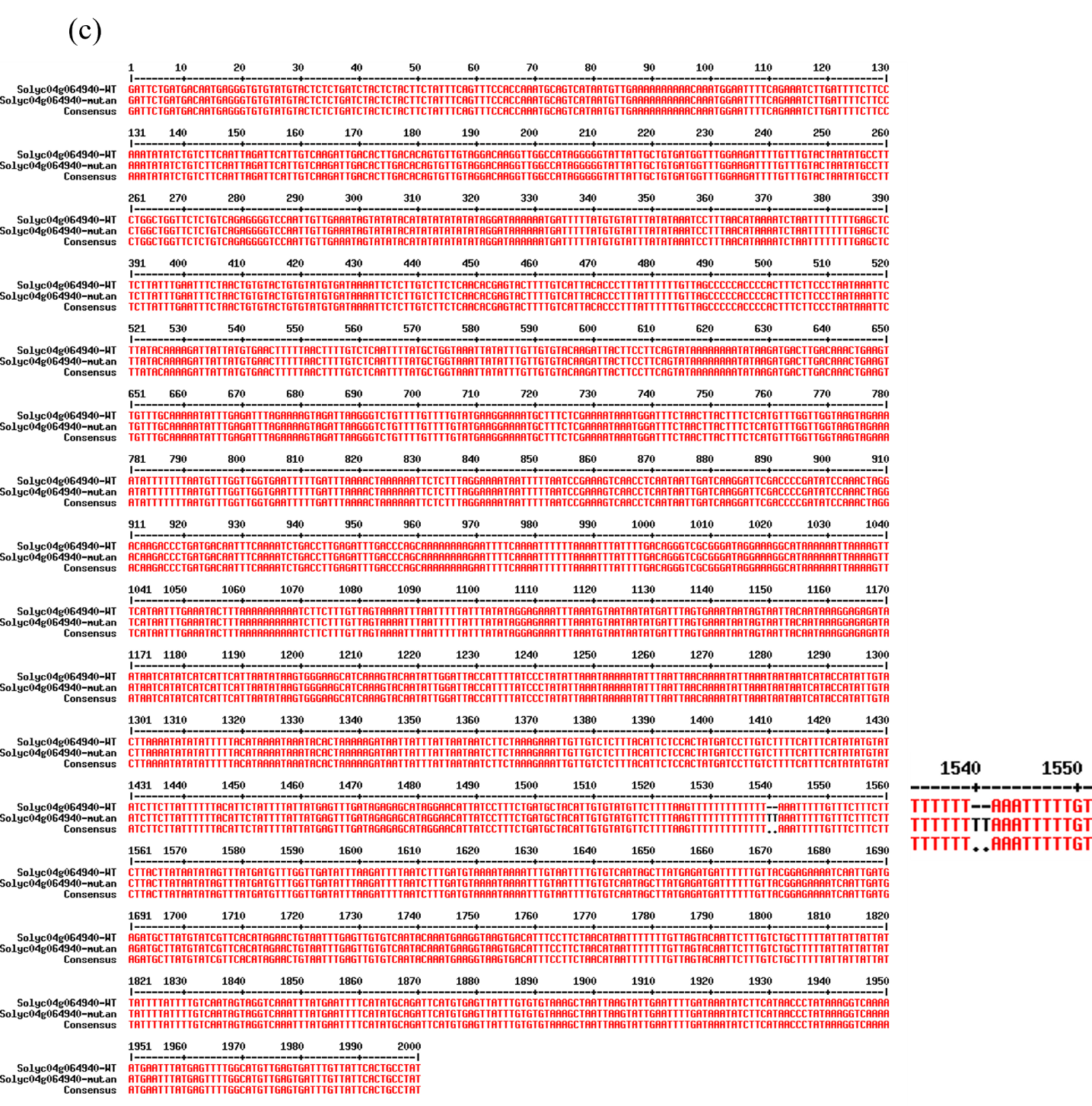
