## Supplementary material for "Root-Suppressed Phenotype of Tomato *Rs* Mutant is Seemingly Related to Expression of Root-Meristem-Specific Sulfotransferases": binding sites of several transcription factors and, in parallel, added binding sites for additional TFs

**Table S2.** Loss or gain of transcription factor binding sites due to promoter mutations. The ~2-kb promoter regions of the sulfotransferase genes *Solyc04g028380*, *Solyc04g028390* and root growth factor receptor kinase gene *Solyc04064940* were analyzed using the Plant ChIP-seq Database o**f *Arabidopsis thaliana* and *Solanum lycopersicum* (PCBase;** <https://pcbase.itps.ncku.edu.tw/promoter_analysis.php>) to predict transcription factor and DNA-binding protein sites. A similarity score threshold of >0.8 was applied, with 1.0 representing an exact match. The table lists transcription factor binding sites altered by the identified promoter mutations. For details, see **Dataset S6**.

| **Gene** | **Mutated base position on promoter** | **Binding site in WT** | **Binding sites in mutant** | **Remarks** |
| --- | --- | --- | --- | --- |
| **Solyc04g028380** | **A-393T** | 1. Transcriptional Repressor of EIN3-dependent Ethylene-response 1 (TREE1) | 1. Transcriptional Repressor of EIN3-dependent Ethylene-response 1 (TREE1)  2. ABF1 | Gained binding site for ABF1 |
| **Solyc04g028390** | **-530**  **T insertion** | SUPPRESSOR OF PHYTOCHROME B4-#3 (SOB3) | MYB3R4  ABF4  DNAJ2  SUPPRESSOR OF PHYTOCHROME B4-#3 (SOB3) | Gained binding sites for  MYB3R4  ABF4  DNAJ2 |
|  | **A-1401T** | SOC1  ABF4  DREB2A  SEP3  FLM  HB5  SOC1  WRKY33  PHERES1 (PHE1)  DREB2A | SOC1  MYB44  SVP  val2  SEP3  FLM  HB5  SOC1  TREE1  WRKY33  PHERES1 (PHE1)  DREB2A | Lost site:  ABF4  Gained binding sites for  MYB44  SVP  val2  TREE1 |
|  | **-1403**  **-1404**  **AT insertion** | MYB3R4  MYB44  HAT22  TREE1 | MYB3R4  SPT6-like (SPT6L)  RGA  PIF4  ELF3  MYB44  MYC2  WRKYs  HAT22  TREE1 | Gained binding sites for  SPT6-like (SPT6L)  RGA  PIF4  ELF3  MYC2  WRKYs |
|  | **-1576**  **A insertion** | B3  (AT3G18990) | B3  (AT3G18990,  AT1G49480) | Same class B3 but gained site for AT1G49480 |
| **Solyc04g064940** | **-1540, -1541**  **TT insertion** | MYB3R4  HB7  MYC3  MYB44  MYC2  WRKY33  DREB2A  HSFA1A  RGA  MYB3R1  WRKYs  TREE1  PIF4  GBF2  BZIP28  val2 | MYB3R4  MYC3  MYB44  MYC2  WRKY33  DREB2A  HSFA1A  RGA  MYB3R1  WRKYs  TREE1  PIF4  GBF2  val2  SPT6L  SEP3  ABF1  HAT22 | Lost site:  HB7  BZIP28  Gained binding site for  SPT6L  SEP3  ABF1  HAT22 |
