## Supplementary material for "Root-Suppressed Phenotype of Tomato *Rs* Mutant is Seemingly Related to Expression of Root-Meristem-Specific Sulfotransferases": the presence of three secondary mutations, which may also influence the observed phenotypes

**Table S3.** The accession and mutant gene details of LA1796. The details were collated from the TGRC in Davis, California, USA. The *Rs* mutant also possesses three additional mutations, namely *d*, *h,* and *gs* ([https://tgrc-mvc.plantsciences.ucdavis.edu](https://tgrc-mvc.plantsciences.ucdavis.edu/Genes/Search))*.*

**Accession Details**

| **Accession** | LA 1796 |
| --- | --- |
| **Status** | Active |
| **Taxon (Solanum)** | *S. lycopersicum* |
| **Formerly** | *L. esculentum* |
| **Donor** | Ernie Kerr |
| **Background genotype** | Hybrid |
| **Mating System** | Autogamous-SC |
| **Horticultural Recommendation** | Graft onto nonmutant rootstock |
| **Categories** | Misc markers; Monogenic |
| **Year accessioned** | 1977 |
| **Genes** | *d, gs, h, Rs* |

**Gene Details**

| **Gene** | **Locus Name** | **Chromosome** | **Gene Sequence** | **Reference** |
| --- | --- | --- | --- | --- |
| *Rs* | Root suppressed | 4 | Not known | **Kerr** (1972) TGRC reports. **22:** 12 |
| *d* | dwarf | 2 | Brassinosteroid C-6 Oxidase (CYP85A1) (Solyc02g089160).  The mutation affects the canonical splice donor site at the junction of exon five and intron 5 | **Nomura et al.** (2005) *Proc. Nat. Acad. Sci.* **102**: 2564–2569. |
| *gs* | green stripe | 7 | TAGL (Solyc07g055920)  Methylation of the TAGL1 promoter, linked to a SNP at SL2.50ch07_63842838 | **Liu et al**. (2020) *New Phytologist*. **228:** 302-17. |
| *h* | hairs absent | 10 | C2H2-type domain-containing protein (Solyc10g078970)  The mutation results from a large chromosomal deletion (1432 bp) in the gene. | **Chang et al.** (2018) *Plant Journal.* **96**: 90-102. |

**Phenotype Details**

| **Allele** | **Source** | **Reference** | **Phenotype** |
| --- | --- | --- | --- |
| *Rs* | Radiation | **Yu & Yeager** (1960) *JASHS* **76**:538. In cv. Chatham. | Greatly restricted or no root development. |
| *d* | Spontaneous | **Butler** (1952) *J Hered* **43**:25 | Shortened hypocotyl, darker, broader, and shorter cotyledons; stems heavy and erect; plant compact, internodes shortened to about 2.5 cm; leaves very distinct, with reduced number and size of segments; dark green color, puckered rugose surface, down-curled margins, and broader, shorter outline of whole as well as individual segments; similar reduction in size and foreshortening of inflorescence, flowers, and fruit. |
| *gs* | Spontaneous | **Larson & Pollack** (1951) *TGC* **1**:9. | Irregular longitudinal green stripes in the epidermis of unripe fruit; retaining chlorophyll for a longer period during ripening, and eventually assuming a paler colour in fully ripe fruit; changes are limited to the epidermis. Striping may be observed on the stem under conditions of high humidity and low light; heterozygotes exhibit some faint striping on the fruit, but are scored as a recessive. |
| *h* | Spontaneous | **Butler (**1952) *J Hered* **43**:25. | Long trichomes are absent except on the hypocotyl and at the growing point; the heterozygote is intermediate. |
